# Rapid characterisation of hERG channel kinetics II: temperature dependence

**DOI:** 10.1101/609719

**Authors:** Chon Lok Lei, Michael Clerx, Kylie A. Beattie, Dario Melgari, Jules C. Hancox, David J. Gavaghan, Liudmila Polonchuk, Ken Wang, Gary R. Mirams

## Abstract

Ion channel behaviour can depend strongly on temperature, with faster kinetics at physiological temperatures leading to considerable changes in currents relative to room temperature. These temperature-dependent changes in voltage-dependent ion channel kinetics (rates of opening, closing and inactivating) are commonly represented with Q_10_ coefficients or an Eyring relationship. In this paper we assess the validity of these representations by characterising channel kinetics at multiple temperatures. We focus on the hERG channel, which is important in drug safety assessment and commonly screened at room temperature, so that results require extrapolation to physiological temperature. In Part I of this study we established a reliable method for high-throughput characterisation of hERG1a (Kv11.1) kinetics, using a 15 second information-rich optimised protocol. In this Part II, we use this protocol to study the temperature dependence of hERG kinetics using CHO cells over-expressing hERG1a on the Nanion SyncroPatch 384PE, a 384-well automated patch clamp platform, with temperature control. We characterise the temperature dependence of hERG gating by fitting the parameters of a mathematical model of hERG kinetics to data obtained at five distinct temperatures between 25 and 37 °C, and validate the models using different protocols. Our models reveal that activation is far more temperature sensitive than inactivation, and we observe that the temperature dependency of the kinetic parameters is not represented well by Q_10_ coefficients: it broadly follows a generalised, but not the standardly-used, Eyring relationship. We also demonstrate that experimental estimations of Q_10_ coefficients are protocol-dependent. Our results show that a direct fit using our 15 second protocol best represents hERG kinetics at any given temperature, and suggests that predictions from the Generalised Eyring theory may be preferentially used if no experimentally-derived data are available.

**Statement of Significance:** Ion channel currents are highly sensitive to temperature changes. Yet because many experiments are performed more easily at room temperature, it is common to extrapolate findings to physiological temperatures through the use of Q_10_ coefficients or Eyring rate theory. By applying short, information-rich protocols that we developed in Part I of this study we identify how kinetic parameters change over temperature. We find that the commonly-used Q_10_ and Eyring formulations are incapable of describing the parameters’ temperature dependence, a more Generalised Eyring relationship works well, but remeasuring kinetics and refitting a model is optimal. The findings have implications for the accuracy of the many applications of Q_10_ coefficients in electrophysiology, and suggest that care is needed to avoid misleading extrapolations in their many scientific and industrial pharmaceutical applications.

## INTRODUCTION

Ion channel behaviour can depend strongly on temperature (1, 2), with physiological temperatures typically leading to faster kinetics and different magnitudes of current than at room temperature, see for example (3, Figure 1). These temperature-dependent changes in voltage-dependent ion channel kinetics, e.g. rates of activation, deactivation, inactivation and recovery, are commonly represented with either Q_10_ coefficients or an Eyring relationship. However, a detailed comparison between these different representations for ion current modelling has not yet been undertaken. Here we characterise channel kinetics at multiple temperatures, and test the validity of Q_10_ and Eyring rate theories by testing whether the kinetic parameters follow the trends that these theories assume. For this case study we use the hERG channel which has been shown to have temperature-dependent kinetics (3–5).

**Figure 1:**
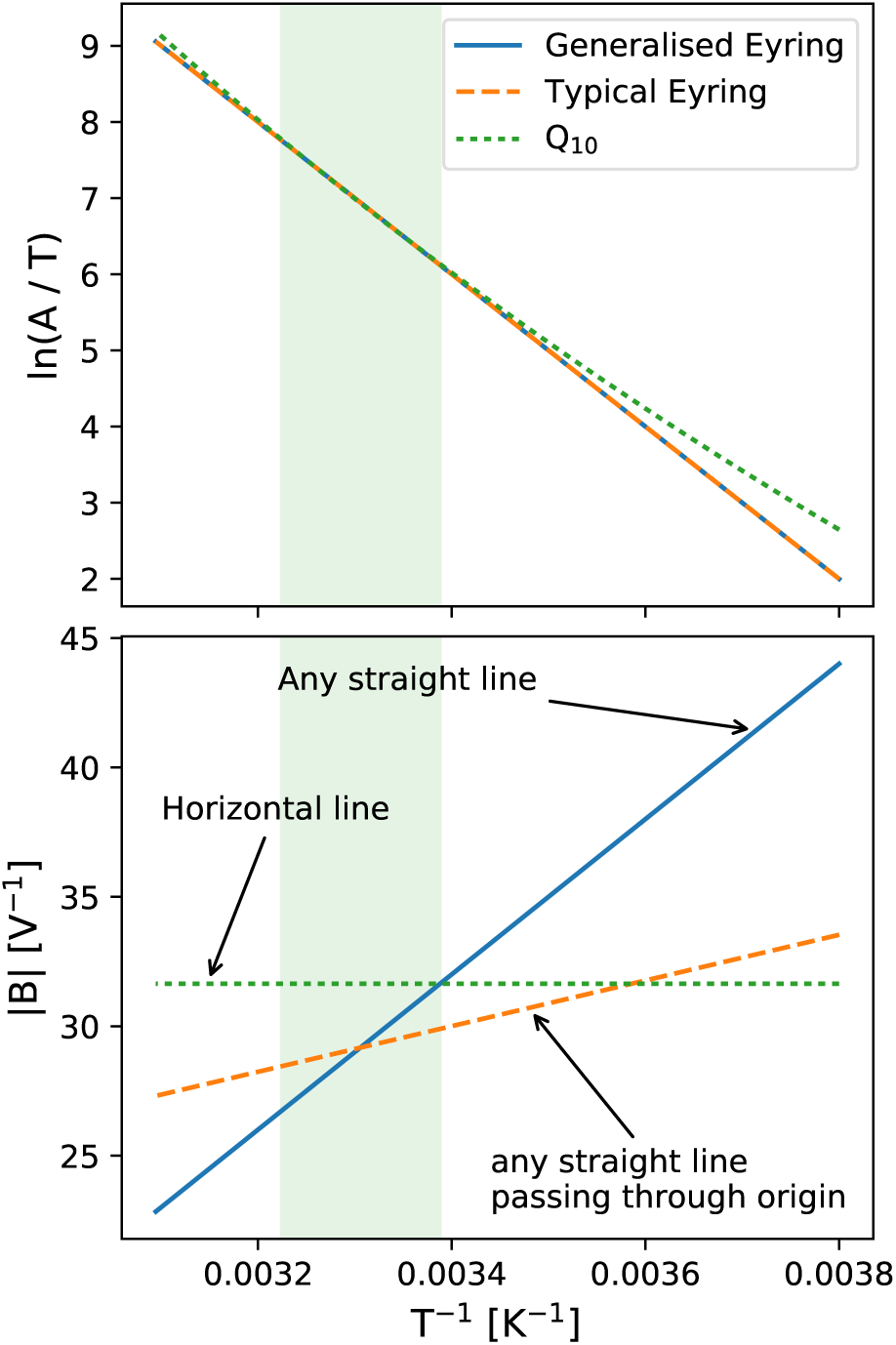
An Eyring plot illustrating the difference between a Generalised Eyring equation (Eq. 4), a Typical Eyring equation (Eq. 3), and a Q_10_ formulation (Eq. 9). This plot extends from −10 °C to 50 °C to highlight the differences between the three formulations. The green shaded region marks the temperature range of interest, from 22–37 °C. The Generalised Eyring relationship shown has [ln *a*_GE_, *b*_GE_, *c*_GE_, *d*_GE_] = [40, 1000, 3000, −70], and the Typical Eyring and Q_10_ relationships are the best fits to the generated Generalised Eyring relationship. Both Eyring formulations give the same straight line dependence for ln(*A*/*T)* and even the nonlinear Q_10_ formulation is indistinguishable (for practical purposes) within the relevant temperature range. However, the three formulations can display very different behaviour when examining the temperature dependence of the voltage-dependence parameter *B*.

The *human Ether-à-go-go-Related Gene* (hERG) encodes the pore-forming alpha subunit of the ion channel Kv11.1 that conducts the rapidly activating cardiac delayed rectifier potassium current (*I*_Kr_) (6). Unless otherwise specified, we refer to hERG1a simply as ‘hERG’ in the remainder of this paper. Pharmaceutical compounds that block *I*_Kr_ can prolong the cardiac ventricular action potential (7), and are associated with both increased QT intervals on the body-surface electrocardiogram and elevated risk of Torsade de Pointes (TdP) arrhythmia in patients (8). The existing International Council for Harmonisation S7B regulatory guidelines for pharmaceutical development require the evaluation of drug effects on the hERG channel as part of pre-clinical safety testing during drug development (9).

Drug effects on hERG are typically characterised by the concentration at which hERG channel activity is reduced by 50% (the ‘IC_50_’) (10). However, no single measurement temperature nor method is used consistently across different laboratories for measuring hERG IC_50_ values. Zhou et al. (4) and Vandenberg et al. (3) measured hERG1a temperature dependence, and compared room and physiological temperature kinetics under typical activation and inactivation current-voltage (I-V) protocols. A similar study with hERG1a/1b was performed more recently by Mauerhöfer and Bauer (5). These studies consistently report that hERG kinetics are highly temperature-sensitive, something that is perhaps a property of potassium channels more widely (2). The use of different temperatures and voltage protocols is therefore thought to be a large source of (deterministic) variation in IC_50_ values (11–13).

In addition, drug screening data is often collected at room temperature, and requires extrapolation to physiological temperature. The temperature extrapolation relies heavily on the accuracy of models of temperature-dependence. Some effort has been made to model temperature effects on hERG kinetics based upon literature data (3, 4); for example Fink et al. (14) attempted to use an Eyring relationship and Li et al. (15) used Q_10_ coefficients. However, a detailed comparison and assessment of the applicability of these representations has not yet been undertaken.

In this paper, we study and model the temperature dependence of hERG kinetics using a cell-specific fitting technique, for a range of room-to-physiological temperatures. We employ a staircase protocol that is applicable in automated high-throughput patch clamp systems, developed in Part I of this study (16). We use a mechanistic model and its parameterisation to characterise hERG kinetics at multiple temperatures, and compare whether these follow the temperature dependence of rate theories. Below, we compare commonly-used temperature adjustments/models for hERG kinetic rates — the Eyring relationship and the Q_10_ coefficient.

### Models of transition rates and their temperature dependence

Mathematical ion channel models are often expressed as a Hodgkin-Huxley model (17) or a Markov state model (18), and both have rates (which we will call *k*) for transitions between the channel gates/states. To derive the rate *k* of transition between two states, the occupancy of two states *p*(*a*), *p*(*b*) at equilibrium is assumed to follow a Maxwell-Boltzmann distribution:

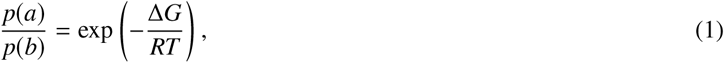

where Δ*G* is the Gibbs free energy difference between the *a* and *b* states, *R* the ideal gas constant, and *T* the absolute temperature. The Gibbs free energy Δ*G* is assumed to be linearly proportional to the membrane potential *V*. Assuming a simple energy barrier model, where only one rate-limiting step is required to transition between two states, the transition rate *k* is then directly proportional to the fraction of system in the excited state, which leads to the commonly-used exponential form (19–21)

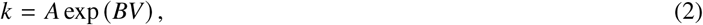

where *A, B* are model parameters (constants). In this study, we use the terms ‘Parameter *A*’ and ‘Parameter *B*’ to refer to *A* and *B* in Eq. 2.

#### Eyring formulations

The temperature dependence of channel transitions is embodied in the Eyring equation. The original Eyring equation was derived from basic thermodynamics and statistical mechanics, following from the the concepts of Gibbs free energy, entropy and enthalpy (22). The typical form used to model voltage-dependent transition rates previously (14, 19, 21, 23) is

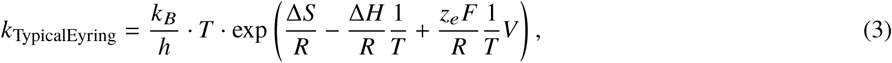

with physical constants: *k*_*B*_ the Boltzmann constant, *R* the ideal gas constant, *h* the Planck constant, *F* the Faraday constant, *T* the absolute temperature, and *V* the trans-membrane voltage. The following are unknowns (or ‘kinetic parameters’) to be determined: Δ*S* the entropy difference, Δ*H* the enthalpy difference, *z*_*e*_ the effective valency of the structure undergoing conformational change. A more generalised Eyring relationship can be given by

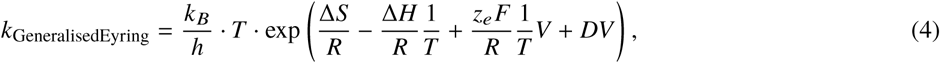

where *D* is a coefficient that describes a temperature-independent effect of voltage on the transition rate. The Generalised Eyring relationship is commonly used in the field of engineering, for example (24–27), although to the best of our knowledge it has not been directly applied to ion channel modelling.

Without loss of generality, we can rewrite (reparameterise) Eq. 4, using unknowns *a*_GE_, *b*_GE_, *c*_GE_, *d*_GE_, absorbing all other constants into these four new parameters, as

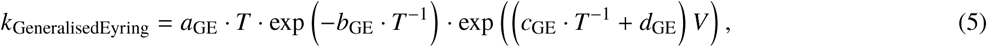

where *a*_GE_ = (*k*_*B*_/*h*) exp(Δ*S*/*R*), *b*_GE_ = Δ*H*/*R, c*_GE_ = (*z*_*e*_ *F*)/*R*, and *d*_GE_ = *D*. By comparing Eq. 2 and Eq. 5, then we have

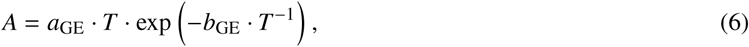

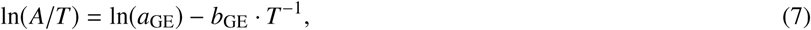

and

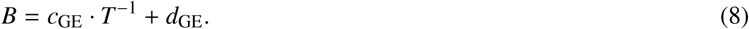

Therefore, plotting ln(*A*/*T*) against *T* ^−1^ should yield a linear relationship if the Generalised Eyring relationship holds. Similarly, from Eq. 8, we see that plotting *B* against *T* ^−1^ yields a linear relationship for the Generalised Eyring relationship; or a proportional relationship for the Typical Eyring relationship (*d*_GE_ = 0). We refer to a plot of ln(*A*/*T*) or *B* as a function of *T* ^−1^ as an ‘*Eyring plot*’.

#### Q_10_ coefficients

Another approach that is commonly used to describe temperature dependence in biological and chemical processes is the use of Q_10_ coefficients. The Q_10_ relationship is an empirical expression (28), which assumes reaction rate increases exponentially with temperature, and has been applied extensively to ion channel kinetics from Hodgkin & Huxley’s work to the present day (3–5, 15, 29, 30). Using Q_10_ coefficients, we can express rates as

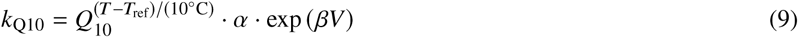

Here, *α* and *β* are parameters for the rate, and *T*_ref_ is the reference temperature for the extrapolation. A *Q*_10_ coefficient is, by definition, calculated using the ratio of the rates at *T*_ref_ + 10°C and *T*_ref_. Comparing Eq. 2 and Eq. 9, we have

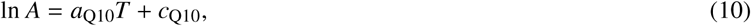

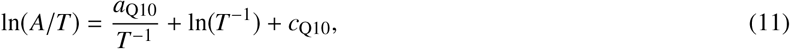

and

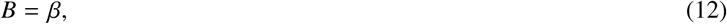

where *a*_Q10_ = (ln *Q*_10_)/10°C, *c*_Q10_ = ln *α* − (*T*_ref_ ln *Q*_10_)/10°C. Therefore, if the Q_10_ formulation is accurate, then plotting ln(*A*/*T*) against *T* ^−1^ should yield a non-linear relationship, and *B* against *T* ^−1^ is a horizontal line.

### A theoretical comparison of the Eyring formulation and Q_10_ coefficient

We now compare the Generalised Eyring relationship (Eq. 4), the Typical Eyring relationship (Eq. 3) and the Q_10_ expression (Eq. 9). Note that the Eyring relationships have been related to the Q_10_ expression (19, 31) to interpret the Q_10_ coefficient as the change of entropy and enthalpy. However, in this study, we treat the two formulations independently.

For parameter *A* in Eq. 2, under the Eyring plot which we plot ln(*A*/*T*) (on the *y*-axis) against 1/*T* (*x*-axis), both the Generalised Eyring and Typical Eyring relationships (Eq. 7) give *y* = *mx* + *c* which is a straight line; while the Q_10_ expression (Eq. 11) becomes *y* = *a*/*x* + ln(*x*) + *b* which is not. This difference could be used to tell which theory is correct, but within our temperature regime the Q_10_ expression on the Eyring plot gives a curve that is indistinguishable, in practical terms, from a linear Eyring relationship, as shown in the top of Figure 1.

Therefore the only practically measurable difference between the potential temperature relationships is in *B* parameters (which set the voltage dependence of the transition rate) in Eq. 2. The Generalised Eyring relationship implies that *B* has a *linear relationship* with *T* ^−1^; the Typical Eyring relationship restricts *B* to be *directly proportional* to *T* ^−1^; and under the Q_10_ coefficient formulation, *B* is a *constant* that does not depend on temperature. These differences are illustrated in the bottom panel of Figure 1.

The Typical Eyring relationship is a special case of the Generalised Eyring relationship, and therefore the Typical Eyring relationship would hold if *D* = 0 were to be obtained when fitting the Generalised Eyring relationship: it will become clear later on that this is not the case for our data. We hence consider and compare only the Generalised Eyring relationship and the Q_10_ formulation in the rest of this study.

There have been previous temperature-dependent hERG modelling studies. Fink et al. (14) expressed hERG kinetics using the Typical Eyring relationship (Eq. 3), but its parameters were derived from experimentally estimated Q_10_ values in Vandenberg et al. (3) yielding an incomplete form of the Eyring relationship based on Q_10_ values. Li et al. (15) used a Q_10_ formulation (Eq. 9) to model temperature dependence of hERG kinetics for simplicity, but did not investigate to what extent this captured temperature-dependent changes in the kinetics.

Modelling temperature effects in ion channel kinetics not only has applications in cardiac safety pharmacology, it is also commonly used in action potential modelling more generally. Many cardiac action potential models (32–35) adapted the Mazhari et al. (36) hERG model which used Q_10_ values from Zhou et al. (4) to extrapolate room temperature recordings to physiological temperature. These extrapolations cause considerable changes to rates, often exceeding changes introduced when modelling diseases or other conditions (37). Similarly, the Christé et al. (38) hERG model was based on measurements at room temperature and extrapolated to 37 °C using Q_10_ values from Vandenberg et al. (3). Within action potential models, many other ion current models (such as *I*_Na_, *I*_CaL_, etc.) are also based on experiments performed at different temperatures (39), most of which are then corrected via Q_10_ extrapolations (40–44).

## MATERIALS AND METHODS

The experimental methods, mathematical model of *I*_Kr_, and the *I*_Kr_ model parameter inference methods used in this paper were identical to the methods detailed in our companion paper (16). We provide only a brief outline of these methods, for details please refer to Lei et al. (16). Here we focus on the methods used specifically for studying the temperature dependence of the channel.

### Experimental methods

Whole-cell patch-clamp voltage clamp experiments were performed on Chinese Hamster Ovary (CHO) cells stably transfected with hERG1a (Kv11.1). Measurements were performed using the Nanion SyncroPatch 384PE (Nanion Technologies GmbH, Germany), an automated high-throughput platform in which each run (or chip) is able to measure 384 wells (with one cell per well) simultaneously. The temperature of machine’s ‘cell hotel’ was set to ~15 °C. Single hole chips with medium resistance (Nanion order number #221102) were used. Solutions used in all measurements are provided in Table S1 in the Supporting Material.

A total of of 9 voltage clamp protocols were used, including the staircase protocol (16), an activation current-voltage (I-V) protocol, a steady-state inactivation I-V protocol, a hERG screening protocol, a delayed afterdepolarization (DAD)-like protocol, an early afterdepolarization (EAD)-like protocol, and action potential-like protocols with beating frequencies of 0.5 Hz, 1 Hz and 2 Hz. A schematic of the experimental procedure is shown in Lei et al. (16, Figure 1). The whole sequence of protocols was applied to every well. Details of these protocols can be found in Lei et al. (16, Supporting Material).

Only the staircase protocol was used in fitting (or calibrating) the mathematical model. The fitted models for each cell were then validated by comparing their predictions for the other 8 protocols to the experimental recordings.

#### Temperature control

The SyncroPatch platform has a temperature control unit with software PE384TemperatureControl, which consists of a temperature controller and several temperature monitors placed around the machine compartment. The machine compartment contains all the solutions on stand-by and is where the measurements occurred. However, since the temperature controller consists of a heater with a fan, it better maintains temperatures higher than room temperature than those close to room temperature. Therefore, the lowest temperature we could maintain indefinitely was 25 °C, which is determined by the room temperature (≈ 22 °C) plus heat generated by the machine’s operation (≈ 3 °C), even if the heat controller itself was set to a lower temperature.

To ensure that we recorded the temperature correctly, an external K-Type thermometer was used to ensure the temperature difference between the measuring stage and the machine in-built temperature monitors was ≲ 0.5 °C. Note that the temperature readouts could differ from the temperature set on the controller even after equilibrium, particularly close to room temperature, so we used the thermometer and temperature monitors’ readouts as the true temperature of the experiments. The temperatures of the five experiments were 25, 27, 30, 33, and 37 °C, and the uncertainty of our temperature measurements was estimated to be ±1 °C by comparing the temperature differences at various locations of the compartment.

#### Post-processing experimental data

We performed a series of quality control checks and corrections (in post-processing) to ensure the currents recorded represent only *I*_Kr_. Leak corrections were applied to all measurements to eliminate leak current (16). E-4031 subtraction was applied to remove any native voltage-dependent ion currents that were present in CHO cells besides the overexpressed hERG1a, usually known as endogenous currents. Cells were then selected based on partially automated quality control described in Part I of this paper (16), resulting in *N*_*e*_ = 124, 91, 85, 84, 45 cells being selected for measurements at 25, 27, 30, 33, and 37 °C, respectively; and our 25 °C data were examined in Part I (16).

#### Data visualisation

Each hERG-transfected CHO cell was expected to have a different total conductance, hence giving a different magnitude for the current recording. Therefore normalisation was applied for visual comparison. Note that the validation of model predictions was performed *without* normalisation (a conductance was fitted for each cell individually). To avoid any circular reasoning involved in normalising based on the *g*_Kr_ parameter fit within the models (which, at this point, may or may not vary with temperature), we used an experimental maximum conductance estimate. The experimental estimate is approximated by extrapolating the negative tail current, after the first 40 mV to −120 mV step, back to the time the voltage step occurred, see Figure S1 in the Supporting Material. Note that this normalisation method is imperfect as it relies on a particular gating process (activation gate *a* ≈ 1 at the end of the 40 mV step) which has some dependence on the kinetics we aim to compare, but the 22 °C parameterisation of the model (45) suggests *a* ≈ 1 is a reasonable approximation (even for lower temperatures) at this point in the protocol. However, since this method removes the conductance dependency, it has a benefit over the normalisation-to-a-reference-trace method used in Paper I (16) by preserving the different magnitudes of currents from *different temperatures*.

### Mathematical model

We used the same Hodgkin & Huxley-style structure hERG model described in the complementary paper Lei et al. (16) and Beattie et al. (45). In this model, the current, *I*_Kr_, is modelled with a standard Ohmic expression,

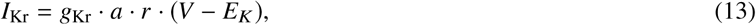

where *g*_Kr_ is the maximal conductance, *a* is a Hodgkin and Huxley (17) activation gate, and *r* is an inactivation gate. *E*_*K*_ is the reversal potential, also known as the Nernst potential, which is not inferred but is calculated directly using

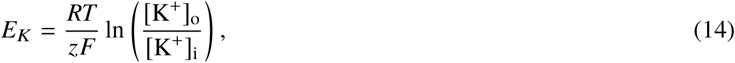

where *R* is the ideal gas constant, *T* is the absolute temperature, *F* is the Faraday constant, and *z* is the valency of the ions (equal to 1 for K^+^). [K^+^]_o_ and [K^+^]_i_ denote the extracellular and intracellular concentrations of K^+^ respectively, which are determined by the experimental solutions, 4 mM and 110 mM respectively. The two gates are governed by

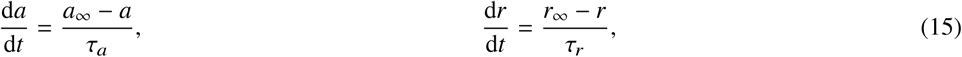

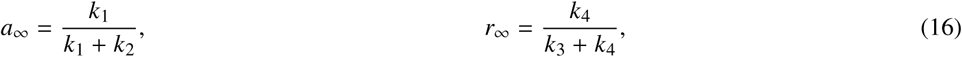

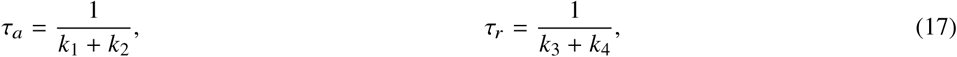

where

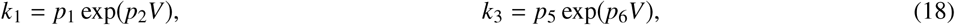

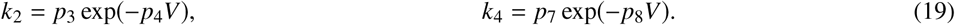

Therefore our model consists of 9 positive parameters ***θ*** = {*p*_1_, …, *p*_8_, *g*_Kr_}, each of is to be inferred from the experimental current recordings.

Simulations were run using Myokit (46), with tolerance settings for the CVODE solver (47) set to `abs_tol` = 10^−8^ and `rel_tol` = 10^−10^. All codes and data are freely available at https://github.com/CardiacModelling/hERGRapidCharacterisation.

### Independent parameter fits at each temperature

The fitting procedure described briefly here follows exactly that laid out in Part I (16), but is repeated for each of the five temperatures.

First, we defined a transformation ***ϕ*** = ln(***θ***) to turn our positively constrained model parameters into unconstrained parameters. For *each temperature*, we specified a statistical model to relate the mathematical model and the observed experimental data:

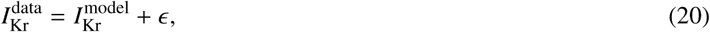

where we assumed the noise term *ϵ* follows a normal distribution *ϵ* ~ 𝒩 (0, *σ*^2^). Writing ***y*** = {*y*_*k*_} for the experimental data 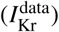 and ***z*** = {*z*_*k*_} for a simulated vector 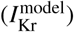, the likelihood of observing a data set **y** given ***ϕ*** is

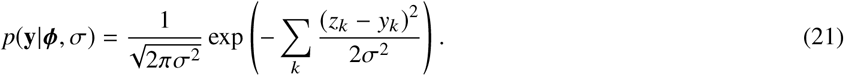

Bayes’ theorem can then be applied to calculate the likelihood of a parameter set given experimental data as

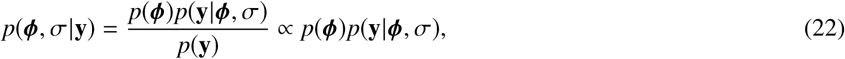

with the prior

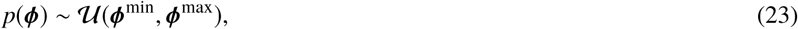

where **𝒰** (·) represents a uniform distribution, for details see Lei et al. (16).

For each temperature *T*, we combined multiple experimental recordings using a hierarchical Bayesian model, as in Lei et al. (16). The full hierarchical Bayesian likelihood is given by

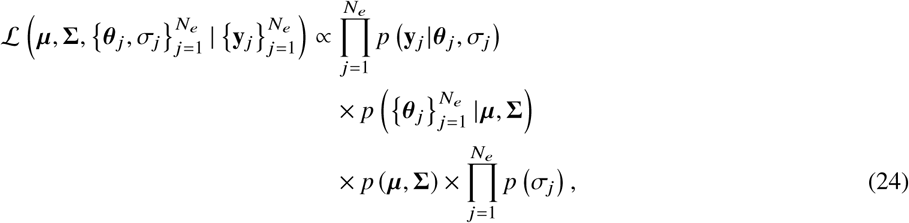

where ***µ*, Σ** are the hyperparameters of the hierarchical model representing the mean vector and covariance matrix from which the individual ‘low-level’ (well-specific) parameters are drawn. 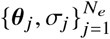 are the set of individual ‘low-level’ parameters for each of the *N*_*e*_ repeats of the experimental recordings 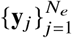. The three terms in Eq. 24 correspond to (a) the likelihood of *all* the individual (low-level) experiments; (b) the likelihood of the hyperparameters (top-level); and (c) the prior of the hyperparameters and the prior of *σ*_*j*_.

We assumed ***ϕ*** _*j*_ for a particular cell (experiment) *j* follows a multivariate normal distribution, namely ***ϕ*** _*j*_ ~ 𝒩 (***µ*, Σ**). There are two distributions describing the well-well variability in this hierarchical Bayesian model, one is the variability of the samples of the mean parameter vector ***µ***, and the other is the covariance matrix **Σ**. As described in the discussion of Lei et al. (16), if we believe the well-well variability represented by **Σ** is primarily due to different patch clamp artefacts in each well, then the uncertainty in ***µ*** represents our uncertainty in the underlying *physiology*, and we therefore believe it corresponds to our uncertainty in the physiological hERG temperature response, rather than our expected variability in the results of future experiments which would require **Σ** too.

For the choice of likelihoods and priors and sampling algorithms we used the simplified pseudo-Metropolis within Gibbs (MwG) algorithm introduced in Part I (16, Supporting Material S6). All inference and sampling were done via our open source Python package, PINTS (48); code is provided as described above.

### Fitting Eyring and Q_10_ relationships

To investigate how well the two temperature models, the Generalised Eyring and the Q_10_ relationships, can explain the temperature dependency of hERG kinetics, we fitted the two temperature models to the inferred distribution of the mean parameter vector ***µ***(*T*) for all temperatures *T*. To do so, first we transformed both the temperature models and ***µ***(*T*) to the Eyring plot form (see Figure 1). Second, we modelled the marginal distribution of ***µ***(*T*) of *p*_*i*_ at each *T* in the Eyring plot using a normal distribution with mean 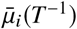 and standard deviation *σ*_*µ,i*_(*T* ^−1^). We further assumed both 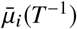 and *σ*_*µ,i*_(*T* ^−1^) follow the temperature models, given by Eq. 7, 8 for the Generalised Eyring relationship and Eq. 11, 12 for the Q_10_ formulation.

Finally, given 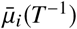 and *σ*_*µ,i*_(*T* ^−1^) (Figure S8 in the Supporting Material), we applied linear regression for parameters *A, B* in the Generalised Eyring model (Eq. 7, 8 to infer *a*_GE_, *b*_GE_, *c*_GE_, *d*_GE_) and a least squares method for only parameters *A* in the Q_10_ relationship (Eq. 11 to infer *a*_Q10_, *c*_Q10_) with the Levenberg–Marquardt algorithm provided in SciPy (49); once to fit the mean and once to fit the standard deviation of each parameter as a function of temperature. Due to the simplicity of the problem after our transformation, a relatively simple optimisation algorithm was sufficient. For the *B* parameter in the Q_10_ relationship (Eq. 12), since it is a constant *β*, we decided to follow the standard way of using the Q_10_ relationship where models are usually extrapolated from room temperature, therefore we directly used 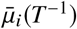 and *σ*_*µ,i*_(*T* ^−1^) at *T* = 25 °C to extrapolate to other temperatures.

The estimated mean as a function of temperature was used to perform predictions for each temperature model; the estimated standard deviation as a function of temperature allowed us to compute the uncertainty bounds of the *I*_Kr_ model parameter for each temperature model.

## RESULTS

### Temperature dependence of recordings

Figure 2 shows the normalised voltage clamp recordings measured with the 9 different protocols, and the corresponding voltage protocols, at the 5 temperatures. For each panel, from top to bottom shows the voltage clamp protocol (black), normalised recordings (blue) that passed quality control at 25, 27, 30, 33, and 37 °C respectively. All results shown are the first of the two repeats of our recordings.

**Figure 2:**
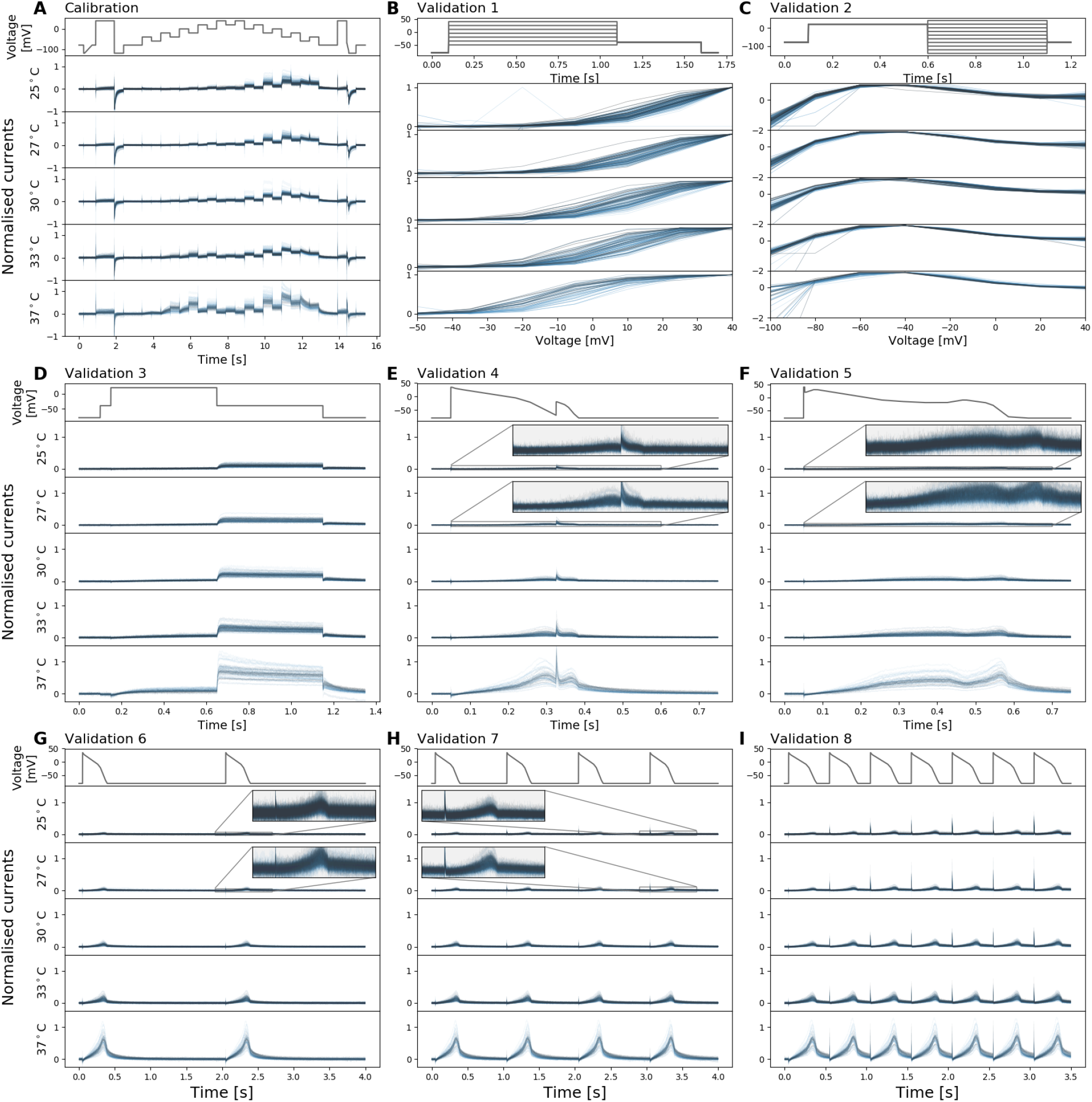
Whole-cell patch-clamp voltage clamp recordings under 9 different protocols, which were all measured in each cell, at 5 temperatures. For each panel, from top to bottom shows the voltage clamp protocol (black), normalised current recordings (blue) that passed quality control at 25, 27, 30, 33, and 37 °C respectively. Currents were normalised with the method described in the text (and see Figure S1). **(A)** Shows the calibration protocol, the staircase protocol. **(B–I)** The eight different protocols used as validation of the model calibration, which are the activation current-voltage (I-V) protocol, the steady-state inactivation I-V protocol, the hERG screening protocol, the DAD-like protocol, the EAD-like protocol, and the cardiac action potential-like protocol at 0.5 Hz, 1 Hz and 2 Hz, respectively. In **(B, C)**, validation 1 and 2 show the I-V relations extracted from the currents.

Figure 2A shows the staircase calibration protocol (in black) and the corresponding experimental recordings (in blue). The change in the recorded current as temperature increased was prominent, it increased the size of the current but also highlighted alterations to the kinetics: during the first half (3–8 s) of the staircase protocol, at low temperature, there was almost no current recorded; however, at physiological temperature, the current was almost as big as the current recorded during the second half (8–13 s) of the staircase protocol. Furthermore, the shape of the current during the second half (8–13 s) of the staircase protocol also changed as temperature increased. This demonstrates that the staircase protocol contains useful information on how kinetics change with temperature.

Figure 2B–I shows experimental recordings for the other 8 validation protocols from the same cells. In validation protocol 1 (Figure 2B) we saw the activation I-V curve shifting to a lower voltage at higher temperatures. In validation protocol 3 (Figure 2D) and validation protocols 6–9 (Figure 2G-I) larger hERG currents were observed at higher temperatures. Both these responses for hERG have been reported previously (3).

### Temperature-dependent fits and predictions

In Lei et al. (16), we showed exclusively the quality of fits and predictions for the hERG models at 25 °C, as this could be most easily compared with previous manual patch results (45); the models replicated both the experimental training *and* validation data very well.

Figure 3 shows the model fitting and validation results for all recorded cells at 37 °C alongside the experimental recordings measured under the 9 different protocols. We fitted the model to the staircase protocol (Figure 3A) and validated against the other 8 protocols (Figure 3B–I). To visually compare the variability in hERG kinetics (and not conductance), currents are normalised by scaling them to minimise the absolute difference between each trace and a reference trace (as in 16). Similar plots for all the intermediate temperatures are shown in the Supporting Material Figures S2–S4.

**Figure 3:**
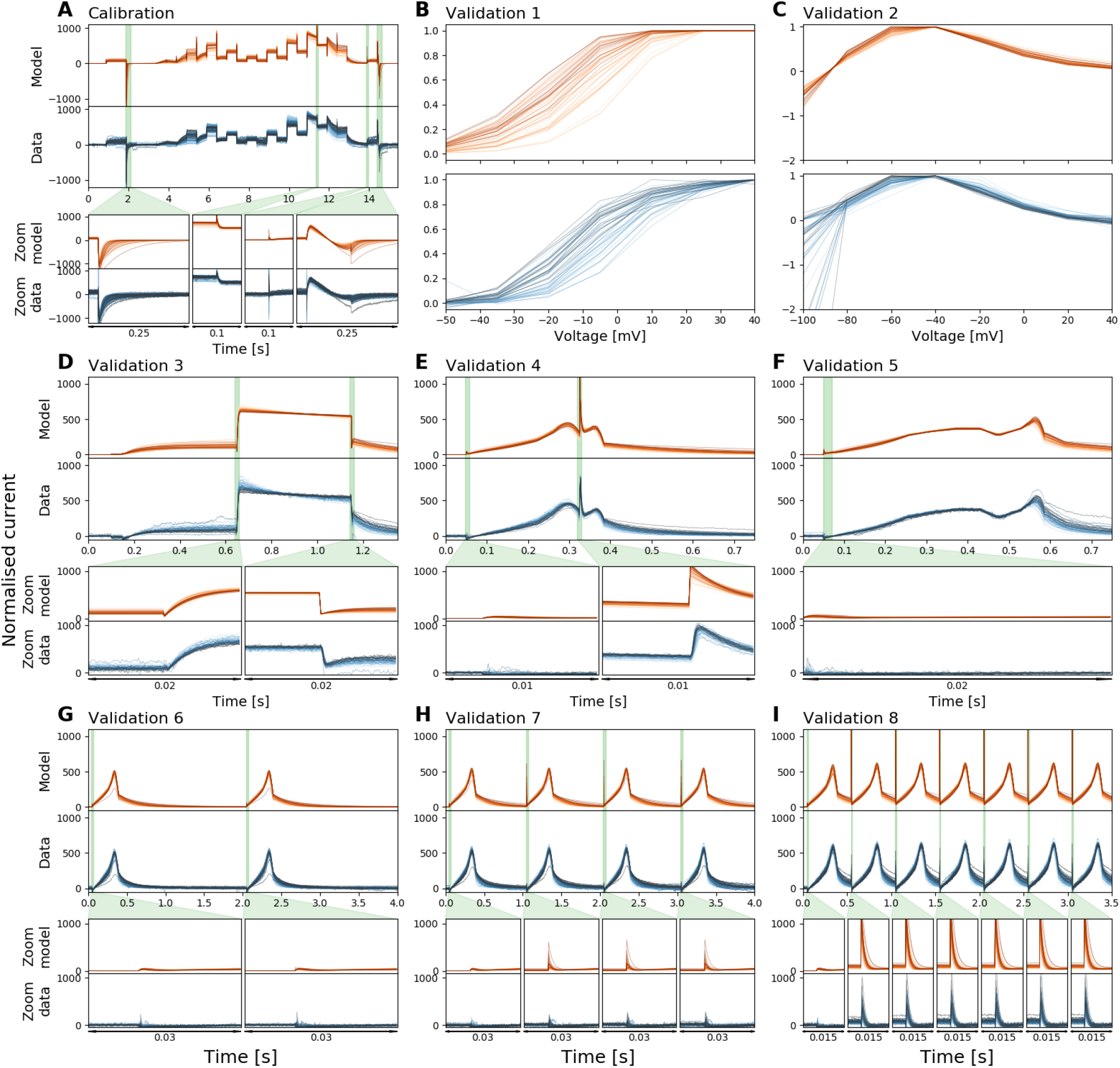
Whole-cell patch-clamp voltage clamp recordings under 9 different protocols, and the model fitting and validation results, at 37 °C. All currents are normalised by scaling them to minimise the absolute difference between each trace and a reference trace. From **(A)** to **(I)**: The staircase protocol which is used as the calibration protocol, the activation current-voltage (I-V) protocol, the steady-state inactivation I-V protocol, the hERG screening protocol, the DAD-like protocol, the EAD-like protocol, and the cardiac action potential-like protocol at 0.5 Hz, 1 Hz and 2 Hz, respectively. All the model calibration results and validation predictions are shown in the top panels (orange), and are compared against the experimental recordings shown in the bottom panels (blue). Zoomed-in of the green shaded regions are shown underneath each panel to reveal the details of the spikes, in which our models show extraordinary good predictions to the details. The normalised current for all protocols are shown except for the activation I-V protocol and the steady-state inactivation I-V protocol where the summary statistic I-V relationships are shown. Each cell is shown with a unique colour.

We applied the same error measure as in Part I of the study to quantify the fits and predictions — the relative root mean square error (RRMSE), defined as

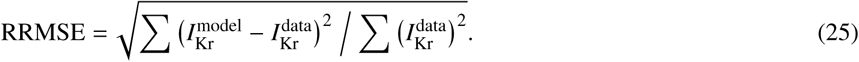

Here, 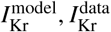 are the model predictions and recordings of *I*_Kr_ respectively. Figure 4 shows the RRMSE histograms for all cells and for the six current trace protocols at 37 °C. Markers indicate the best (*), median (‡) and 90^th^ percentile (#) RRMSE values, and corresponding raw traces and predictions are shown in the three panels above. We note that the models only show single exponential decays due to the limitations of the model structure whilst the data seem to show double exponential decays. These results demonstrate that the hERG model remains a very good representation of the current kinetics, even at 37 °C, the highest temperature. The same analysis has been applied to the intermediate temperatures, the results are shown in Supporting Material Figures S5–S7.

**Figure 4:**
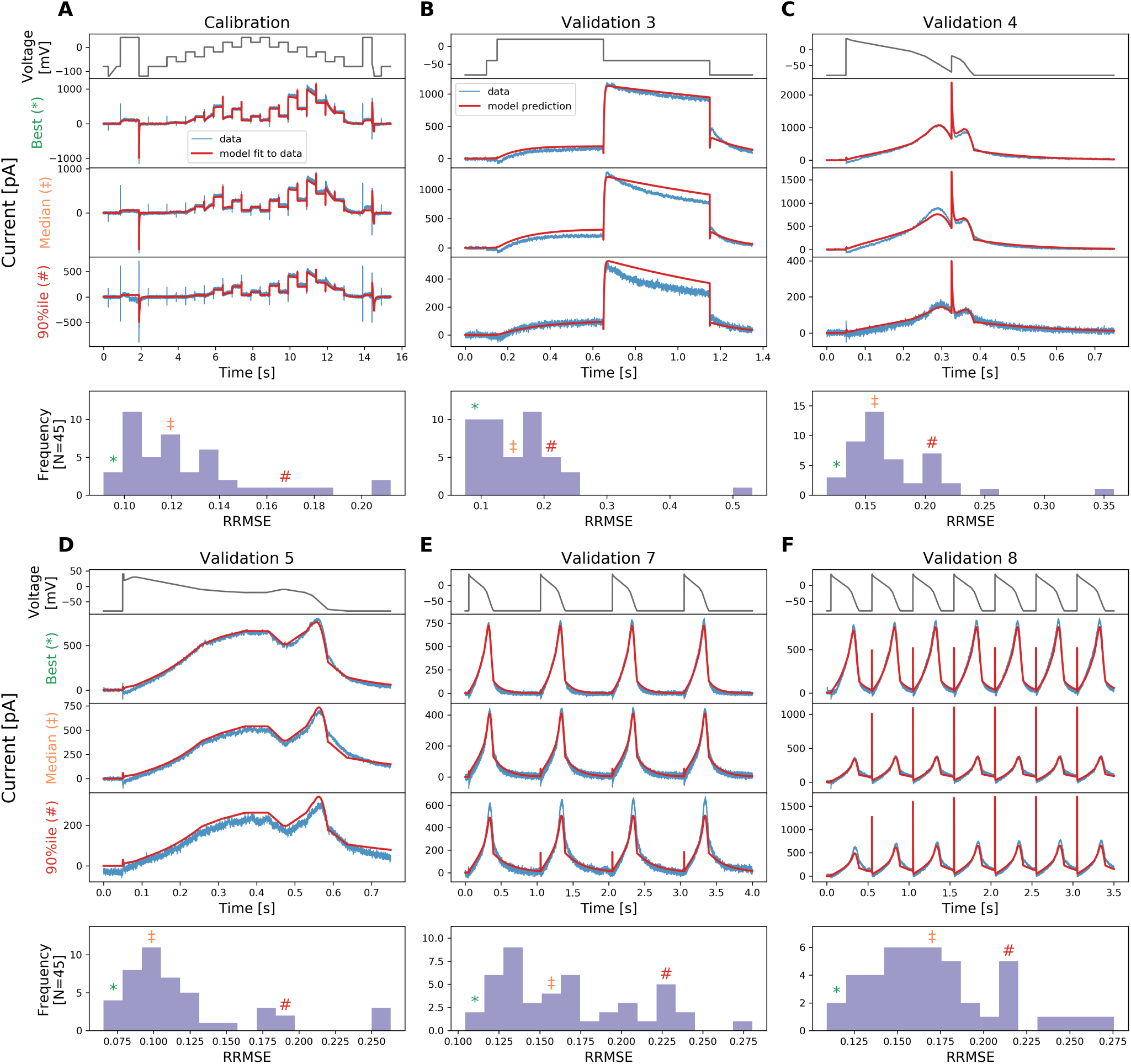
The relative root mean square error (RRMSE, given by Eq. 25) histograms for all cells and for 6 protocols used at 37 °C. Markers indicate the best (*), median (‡) and 90^th^ percentile (#) RRMSE values. The raw traces with the best, median and 90^th^ percentile RRMSE values, for both the model (red) and data (blue), are shown in the panels above, together with the voltage protocol shown on top. Note that the currents are shown on different *y*-axis limits, to reveal the details of the traces.

### Temperature dependence of inferred model parameters

Figure 5 shows the inferred parameter values, which are used in the model predictions in Figure 3 and Figures S2–S4, as a function of temperature. The figure shows the inferred distribution of the hyperparameter mean vector ***µ*** (Eq. 24) using the simplified pseudo-MwG at each temperature in a violin plot.

**Figure 5:**
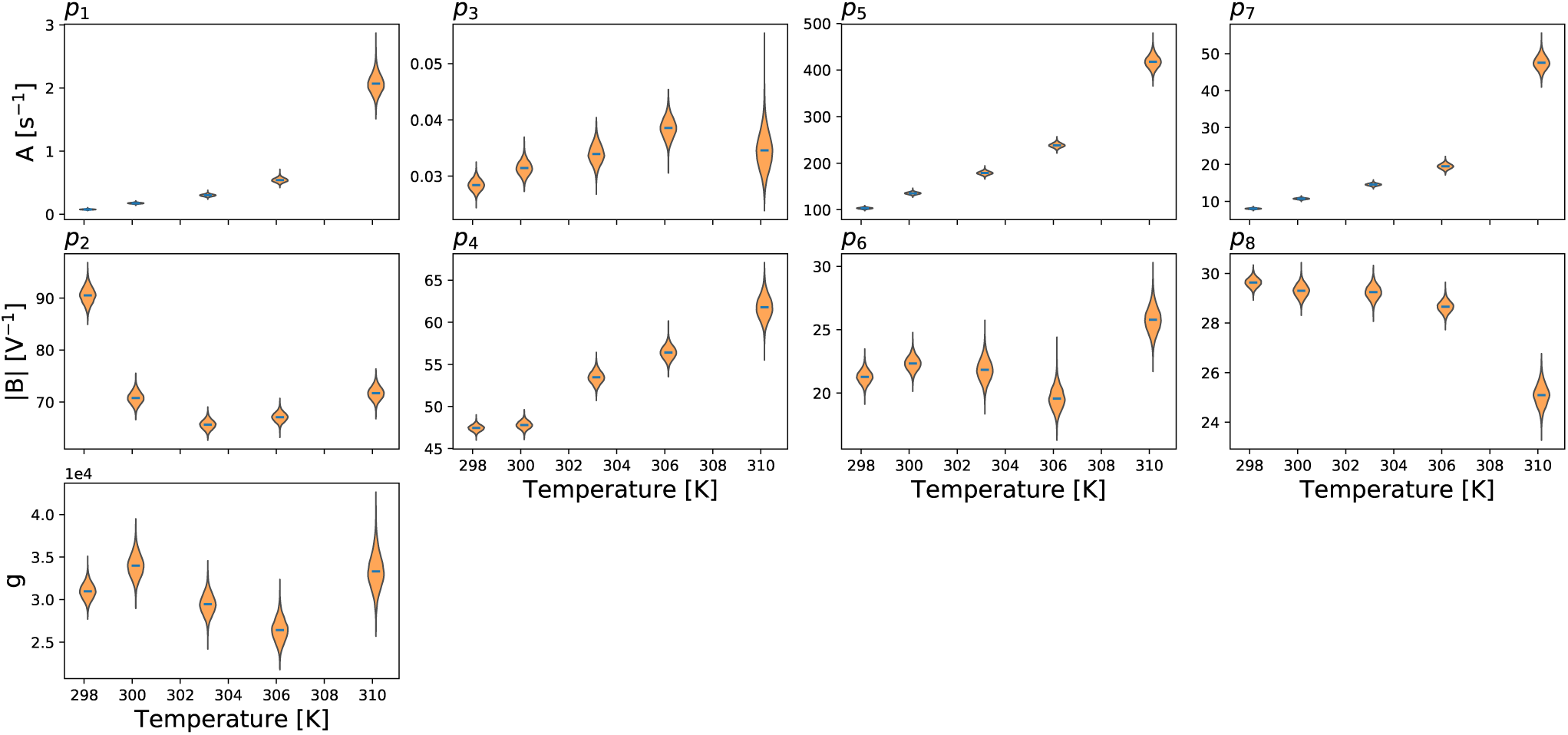
Model parameters plotted as a function of temperature. Here only the inferred distribution of the hyperparameter mean vector ***µ*** (Eq. 24) using the simplified pseudo-MwG at each temperature is shown. Parameters *A, B* refer to Eq. 2. Model parameters show different degrees of temperature dependency. The conductance *g* does not show a prominent change as temperature increases.

If the model kinetics were exhibiting temperature dependence following Q_10_ or Eyring rate theory, then lines whose function is specified by these principles would fit the inferred parameters in Figure 5.

In Figure 5, most parameters show an obvious monotonic trend as temperature increases; although a handful take a slightly more complicated form. It is obvious that the *B* parameters in the second row, *p*_*i*_ with even *i*, are *not* constant over temperatures as would be expected from the Q_10_ relationship. An Eyring plot version of Figure 5 is shown in the Supporting Material Figure S8. We will compare these inferred parameters with the theoretical relationships in detail in the next section.

We then applied Eqs. 16 & 17 to calculate the steady-states *a*_∞_, *r*_∞_ and time constants *τ*_*a*_, *τ*_*r*_ at the 5 temperatures, using the mean of the inferred distribution of ***µ*** at each temperature. Figure 6 shows the resulting voltage dependency of the steady-states and time constants of the model gates *a* and *r*, where each temperature is indicated by a different colour (25 °C blue to 37 °C red).

**Figure 6:**
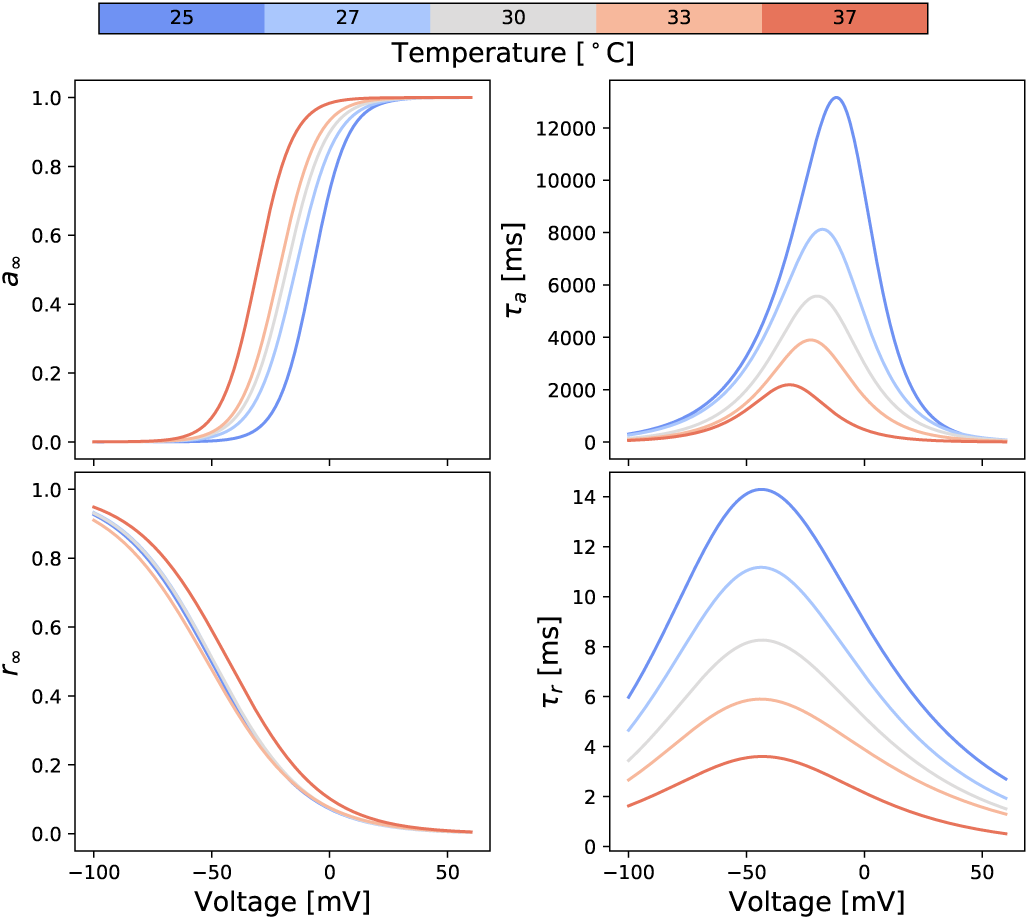
Predicted voltage dependency of steady-states and time constants of the model gates *a* and *r* at different temperatures. These lines are calculated directly from inferred parameters using Eq. 16 & 17 with the independently-fitted hierarchical Bayesian model mean values.

Figure 6 shows that as temperature increases, the steady-state of the activation gate *a* shifts in a negative voltage direction, a prediction from the fitted model that is in agreement with the experimental observations in validation protocol 1: the voltage of half-maximal activation (*V*_1/2_) of *a*_∞_ shifts from −7.5 mV at 25 °C to −30.9 mV at 37 °C, without a noticeable change in the slope factor. However, the steady-state of the inactivation gate *r* does not show a prominent change over temperatures.

The time constant of both gates *τ*_*a*_, *τ*_*r*_ show a similar effect as temperature increases; the maximum *τ*_*a*_ drops from 13.2 s at 25 °C to 2.2 s at 37 °C, and the maximum *τ*_*r*_ drops from 14.3 ms at 25 °C to 3.6 ms at 37 °C. Note that *τ*_*a*_ is in the order of seconds while *τ*_*r*_ is in milliseconds. The voltage which maximises the time constant shifts from −11.6 mV at 25 °C to −31.7 mV at 37 °C for the activation gate, although it does not show a noticeable change for the inactivation gate.

### Comparing models of temperature dependence

Figure 7 shows the Generalised Eyring relationship and the Q_10_ equation fitted to the inferred parameters shown in Figure 5 (orange violin plot). The results are shown in the Eyring plot form: ln(*A*/*T)* and |*B*| as functions of *T* ^-1^. A version of Figure 7 with model parameters plotted directly against temperature is shown in Figure S9 in the Supporting Material. The Generalised Eyring fits are shown as green fan charts with the first three standard deviations; the Q_10_ fits are shown similarly in red. The obtained parameters for the Generalised Eyring equation (Eq. 4) and the Q_10_ equation (Eq. 9) are given in the bottom right tables, one set for each rate *k*_*i*_, *i* = 1, 2, 3, 4. Reassuringly, the values in the tables are comparable to (the same orders of magnitude as) typical literature values for ion channel models.

**Figure 7:**
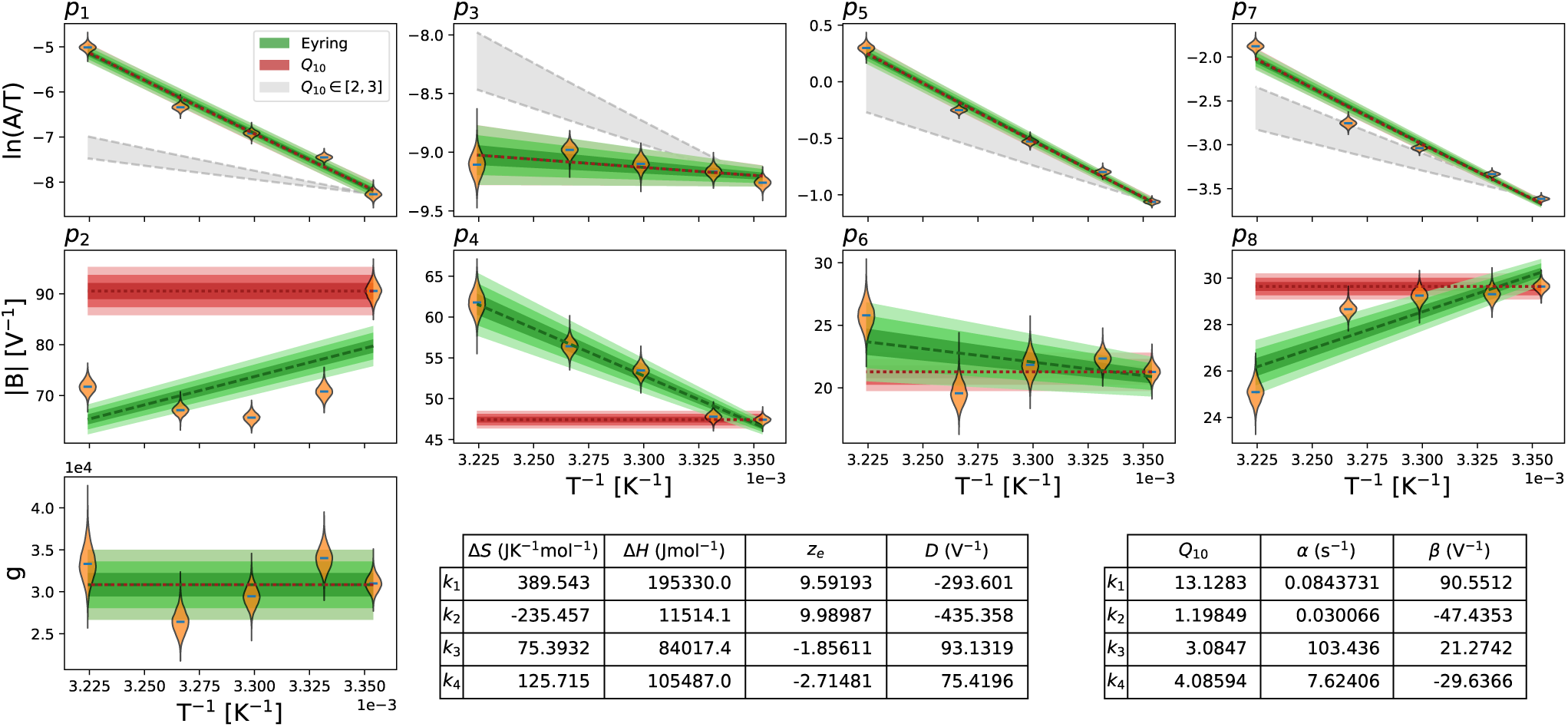
Fitting of Generalised Eyring equation and Q_10_ equation to the distribution of the mean parameter values (mean over all wells, ***µ***, shown with an orange violin plot) on the Eyring axes. The obtained Generalised Eyring fits are shown as green fan charts with the first three standard deviations; the obtained Q_10_ fits are shown in red. The fitted parameters for the Generalised Eyring and Q_10_ equations are shown in the bottom right tables, one set for each *k*_*i*_, *i* = 1, 2, 3, 4. For Q_10_ equations, *T*_ref_ = 298.15 K was used. Note that the non-zero estimations of *D* in the Generalised Eyring relationship indicate that the Typical Eyring cannot fit to all *B* parameters, as it is required to go through the origin. For comparison to typical Q_10_ values in literature, where Q_10_ values are usually assumed to be around 2 to 3, we show the parameters prediction using Q_10_∈[2, 3] as the grey shaded region.

From the illustration in Figure 1, we expect the Generalised Eyring and Q_10_ formulations to be indistinguishable for the *A* parameters, and indeed in Figure 7, the green fan charts (Generalised Eyring) are on top of the red fan charts (Q_10_) in the first row: both formulations are able to fit to the inferred model’s *A* parameters.

Figure 7 shows that the Generalised Eyring equations fit better to the inferred model *B* parameters than the Q_10_ equations. The Generalised Eyring equations are able to fit the inferred model parameters to a large extent except for *p*_2_; whereas the *B* parameters in the Q_10_ equations are not temperature dependent (by definition), which is contradicted by our observations.

Furthermore, it is evident that for parameters *p*_4_, *p*_6_ the two lines cannot intercept the *y*-axis close to the origin, as they are decreasing rather than increasing on these plots. Parameters *p*_2_ and *p*_8_ also have non-zero estimates of *D* in the Generalised Eyring relationship, indicating that the Typical Eyring relationship cannot be fit to any of our *B* parameters. The closest Typical Eyring relationship for *p*_8_ is actually the example shown earlier in the bottom panel of Figure 1. The gradient of the Generalised Eyring fit is approximately twice as steep as the Typical Eyring fit would require for this parameter.

In the literature, Q_10_ coefficients for biological processes such as channel gating are commonly thought to take values from around 2 to 3 (50). To investigate this assumption, we projected our 25 °C model parameters directly using Eq. 9 with Q_10_∈[2, 3] and shown as the grey shaded region in Figure 7. Parameter *p*_5_ in the inactivation rate (*k*_3_) gives a Q_10_ just above 3, but none of our other inferred relationships for parameter *A* is close to the range Q_10_∈[2, 3].

Figure 8 shows the mean model predictions from the temperature-specific parameters (orange), the Generalised Eyring formulation (dotted green), and the Q_10_ coefficient (dashed red) for the staircase protocol. The top panel shows the staircase protocol, followed by the normalised current at 5 different temperatures. Data (in Figure 2A) are shown in fan charts style with the 30^th^, 60^th^ and 90^th^ percentiles in blue. At low temperatures, all three models agree with the data. At higher temperatures, particularly at 37 °C the predictions from the Generalised Eyring formulation (dotted green) still agree reasonably with the temperature-specific independently fitted parameters (orange) and both fit the data (blue) well. However, the prediction from the Q_10_ coefficient deviates from the data during the spikes (see zoomed-in images on the right) and does not predict the time-course accurately during 4–7 s and 12–13 s of the staircase protocol (see insets in Figure 8).

**Figure 8:**
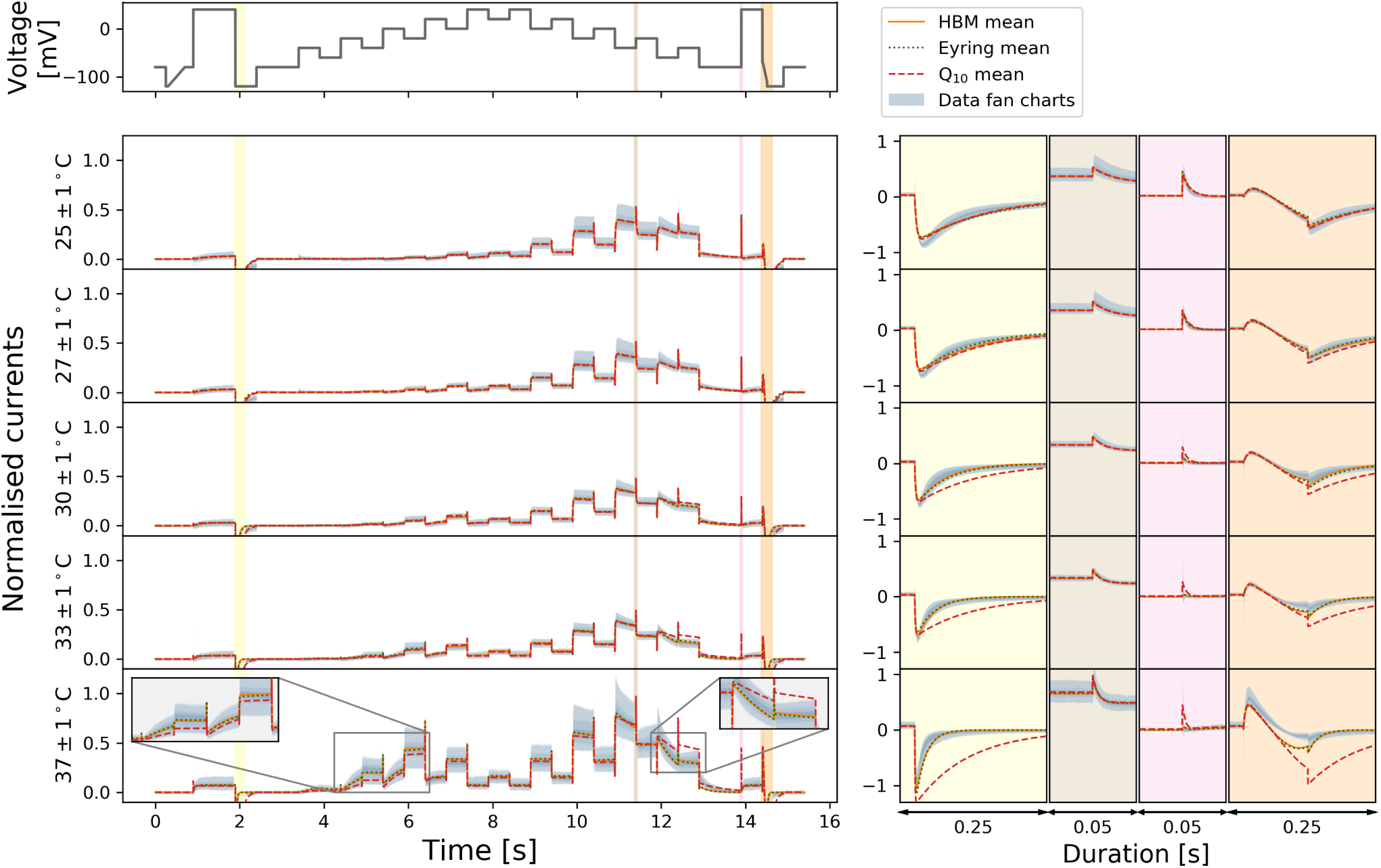
Comparison of the Generalised Eyring formulation (dotted green) and Q_10_ coefficient (dashed red) mean predictions for the staircase protocol. Top figure shows the staircase protocol, followed by the normalised current at 5 different temperatures. Data (in Figure 2A) are shown in fan charts style with the 90^th^, 60^th^ and 30^th^ percentiles in blue. The mean prediction from the hierarchical Bayesian model (HBM) is shown in orange. Zoomed-in to spikes are shown on the right with colours matching to the main plots on the left.

Figure 9 shows a 2 Hz action potential-like protocol prediction version of Figure 8. All the three mean models are able to predict the current during the repolarisation of the action potential clamp very well. The spikes during the upstrokes are however badly predicted by the Q_10_ coefficient mean model; while the Generalised Eyring formulation, similar to the temperature-specific parameters, gives a prediction closer to the data.

**Figure 9:**
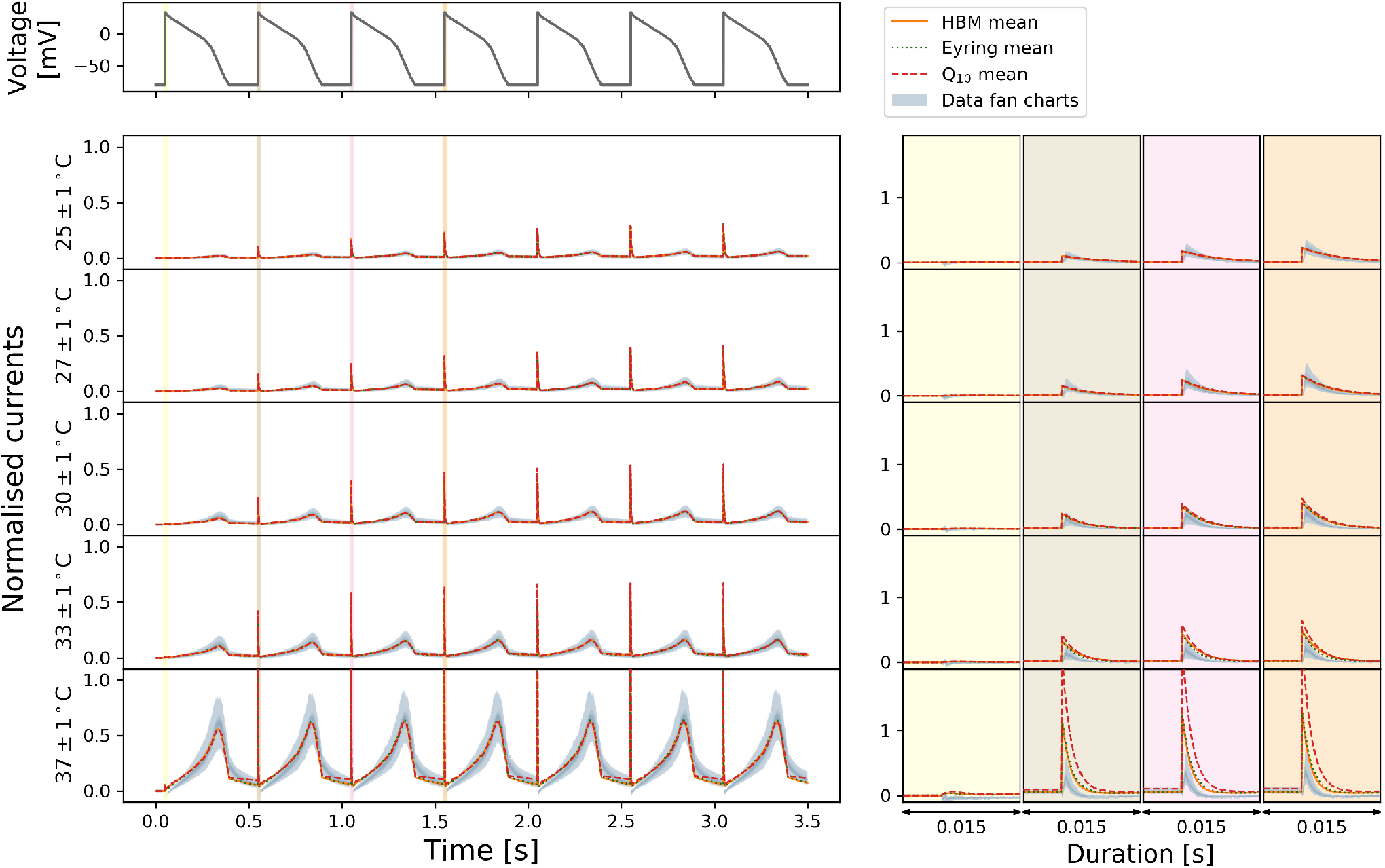
Comparison of the Generalised Eyring formulation (dotted green) and Q_10_ coefficient (dashed red) mean predictions for the 2 Hz action potential-like protocol. Top figure shows the staircase protocol, followed by the normalised current at 5 different temperatures. Data (Figure 2A) are shown in fan charts style with the 90^th^, 60^th^ and 30^th^ percentiles in blue. The mean prediction from the hierarchical Bayesian model (HBM) is shown in orange. Zoomed-in to spikes are shown on the right with colours matching to the main plots on the left.

## DISCUSSION

In this study we have examined the temperature dependence of hERG kinetics, at five temperatures ranging from room to body temperature, with 45 to 124 cells per temperature. We have used a mechanistic model and its parameterisation to capture our knowledge of the hERG kinetics. By assuming that all cells share the same mechanism underlying hERG kinetics we have based our study on the inferred model parameters at different temperatures to reveal the temperature dependence of hERG gating kinetics. This is, to our knowledge, the first systematic effort to have taken this approach.

Using the staircase protocol, we were able to characterise hERG kinetics to the extent that our model can replicate both the experimental training *and* validation data very well, for *all* of the measured temperatures. Our models can predict the current response to the physiologically-relevant action potential protocols with a very high accuracy, demonstrating that our *I*_Kr_ models are robust in predicting hERG current, in both healthy and arrhythmic situations. This gives us confidence that the *cell-specific model* parameters do represent and capture hERG kinetics at the given temperatures.

The directly fitted models reveal that the activation gate has a much higher temperature sensitivity than the inactivation gate. This effect is shown in both the comparison of steady-states and time constants (Figure 6) and the inferred Q_10_ coefficients (Figure 7) where the Q_10_ values for the activation gate (*k*_1_, *k*_2_) are overall higher than the inactivation gate (*k*_3_, *k*_4_). Our inferred Q_10_ coefficient for the rate of activation (*k*_1_) is relatively high compared to literature results (3, 4). However, our findings are not implausible, when compared to other potassium channels, such as Kv2.1 and Kv4.3, which can have maximum Q_10_ values up to the 20–30 range (2). Other ion channels can also exhibit a very high temperature sensitivity, such as TRP ion channels which were reported to have Q_10_ values ranging from 2–15 in Dhaka et al. (1). We then further compare our model predictions with the literature results in Vandenberg et al. (3).

Our hierarchical Bayesian models at different temperatures are not only able to predict our validation data but also able to reproduce the temperature dependence seen in previous studies (3), where the increase of temperature caused a large increase in the overall ‘steady state open probability’. In Section S8 of the Supporting Material, we describe how we reproduced Figure 6 of Vandenberg et al. (3). Figure 10 shows that our simulations (right panel) are broadly consistent with the temperature effect observed in Vandenberg et al. (3) (left panel). The fan charts show the 30^th^, 60^th^ and 90^th^ percentiles of the simulations, representing the *inter-experiment* (well-well) variability. There are differences between our simulations and their experimental results, with a smaller open probability at low temperatures in our simulations and a slight shift of the curves to the right. Nevertheless, our results are broadly consistent with the temperature effect observed in Vandenberg et al. (3) and predict a very similar ‘width’ for this steady state window of open probability and also agree with the absolute values of the probabilities at the higher temperature very well.

**Figure 10:**
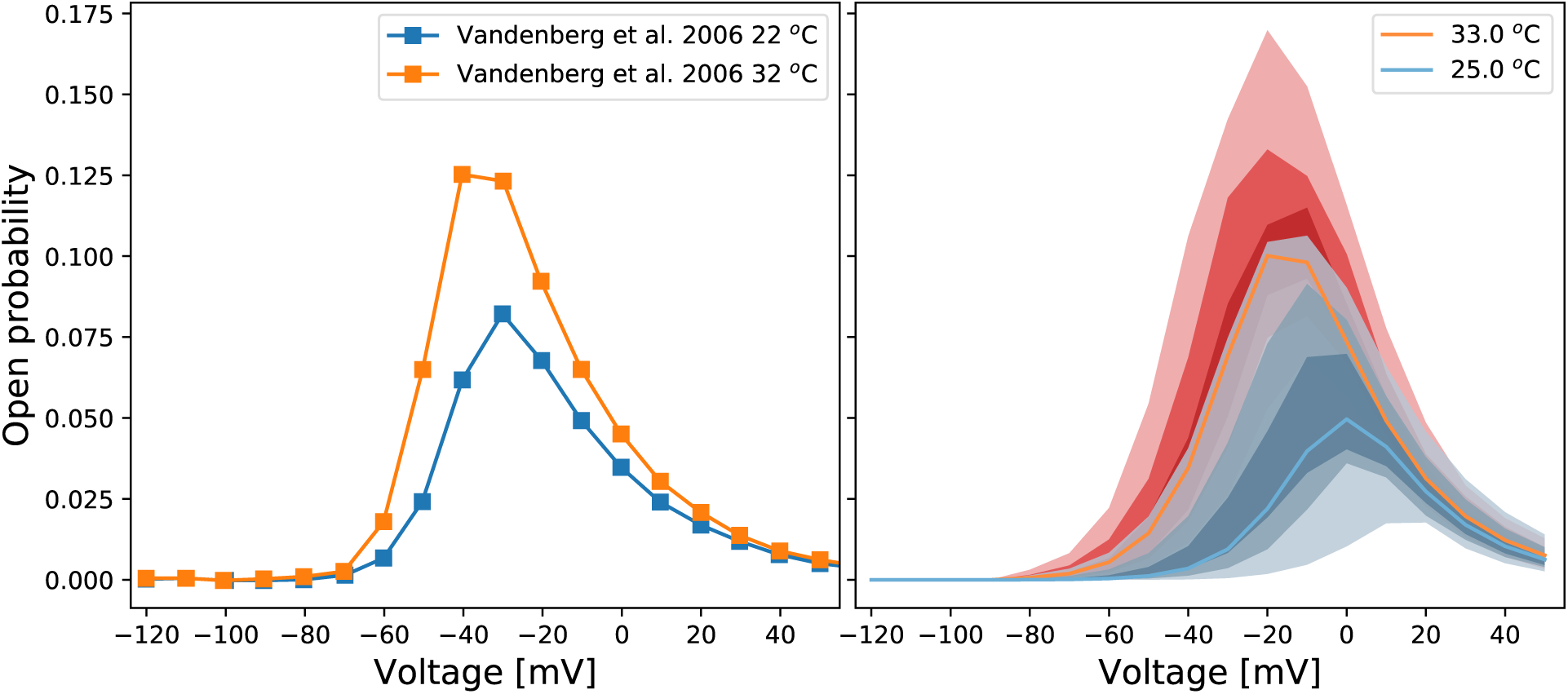
Voltage dependence of steady state ‘open probability’ as defined in Vandenberg et al. (3, Figure 6) using a multiplication of experimental approximations for the product *a*_∞_*r*_∞_. **Left:** Data extracted from Vandenberg et al. (3, Figure 6). **Right:** The fan charts show the the 90^th^, 60^th^ and 30^th^ percentiles of the hierarchical Bayesian model simulations, representing the *experiment-experiment* variability. Orange/red represents 32–33 °C, and blue represents 22–25 °C.

Q_10_ formulations have often been estimated in the past with different protocols, even for the same gating process (e.g. activation). For example, two well-known experimental studies of temperature dependence of hERG kinetics, by Zhou et al. (4) and Vandenberg et al. (3), estimated the Q_10_ coefficients using different protocols and analyses, and reported two very different sets of Q_10_ coefficients (see Table 1) for various gating processes. We asked, if the two experiments were to be repeated with the *same underlying kinetics*, would they agree with one another? Using our directly-fitted models at 25 °C and 37 °C, we simulated the two different sets of experiments described in (3, 4), for details see Supporting Material Section S9. We then estimated two sets of Q_10_ coefficients following the protocols and analysis in each of the papers, and the obtained values are shown in Table 1. The findings in Table 1 show strong evidence that due to *different protocols* the estimated Q_10_ coefficients can disagree. Furthermore, neither of the protocols reproduces the direct estimate of Q_10_ coefficients from the model parameter temperature relationships (shown in the bottom right of Figure 7). We conclude that extreme caution should be used when directly modifying rates in models with experimental estimations of Q_10_ coefficients.

**Table 1:** Q_10_ coefficients for each gating process, estimated using the protocols specified in Zhou et al. (4) and Vandenberg et al. (3). The values were derived from simulations performed using the temperature-specific hierarchical Bayesian model fits at 25 °C and 37 °C.

|  | Zhou et al. (4) |  | Vandenberg et al. (3) |  |
| --- | --- | --- | --- | --- |
|  | Reported values | Model estimation | Reported values | Model estimation |
| Activation | $6.25 \pm 2.55$ | $10.668 \pm 7.482$ | $2.1 \pm 0.30$ | $7.400 \pm 4.111$ |
| Deactivation | $1.36 \pm 0.40$ | $2.016 \pm 0.764$ | $1.7 \pm 0.30$ | $3.692 \pm 1.224$ |
| Inactivation | $3.55 \pm 0.87$ | $3.421 \pm 1.028$ | $2.5 \pm 0.53$ | $2.750 \pm 0.900$ |
| Recovery | $3.65 \pm 0.73$ | $2.991 \pm 0.730$ | $2.6 \pm 0.26$ | $4.436 \pm 2.763$ |

Fitting directly to the staircase protocol at different temperatures does not require any assumption about the underlying temperature dependence of the kinetic parameters, except that the model structure does not change. The existing well-known models/approximations for temperature dependence of ion channel transition rates are the Q_10_ and Typical Eyring formulations. Our study has raised concerns about how accurate these relationships are. In terms of parameter values (Figure 7) none of these methods is able to capture the full temperature dependence of the directly-fitted parameters, ***µ***(*T*), and predictably this impairs their ability to fit and predict currents (Figure 8). However, using a Generalised Eyring relationship (not commonly used in ion channel modelling) can closely mimic our full direct fitting approach (Figure 8, 9). Whilst the model predictions using the Q_10_ formulation can generally predict overall trends in temperature effects, the predictions cannot capture the details of the current, compared to the Generalised Eyring relationship or the full direct fitting approach (Figure 8 and 9). We therefore suggest neither Q_10_ formulations nor the Typical Eyring relationship should be used: the Generalised Eyring relationship is much better for temperature predictions. But for the best results the model should be refitted at any temperature of interest using an information-rich protocol, such as our staircase protocol (16).

The nonlinearity of some kinetic parameters on the Eyring plots implies the Generalised Eyring relationship is a reasonable but imperfect temperature model. Under the assumption that the model structure is correct, we accurately captured the kinetics at each temperature, and the model structure stays the same for all temperatures. However, we could challenge these assumptions, and suppose that the Generalised, or even Typical, Eyring relationship is true for any transition of ion channel from one conformational state to another. In this case, the Eyring formulation not matching the individual temperature parameter sets could imply that, either: (a) the hERG model structure that we have assumed is incorrect, i.e. the relationship not holding is a consequence of discrepancy between the model and reality; or, (b) our procedure did not accurately capture the kinetic parameters at each temperature, but the fact that the parameters give excellent fits and predictions (and many parameters do follow expected trends) perhaps alleviates this concern; or, (c) in reality, the energy landscape of ion channel conformations changes with temperature, and a given transition in the model represents a different jump in conformational state, i.e. the model structure should change with temperature (which has been modelled previously (51)).

In any case, applying a simple treatment such as the Generalised Eyring relationship or the Q_10_ coefficient to an imperfect model that violates the assumptions above would not automatically alleviate any mismatch. Since our temperature-specific fits can replicate both the experimental training data *and* the validation data very well at all temperatures, the model is a good *representation* of hERG kinetics. Hence it is better to apply a rapid and reproducible procedure, as illustrated here, for generating all the parameters within a model at a new temperature, whenever possible. However, if necessary, then the Generalised Eyring relationship would be a preferable choice for predicting kinetics at a new temperature where measurements cannot be, or have not been, taken. While further work might show our results are more generally applicable to other channels, for now they should be interpreted as being specific to hERG1a.

Our results have strong implications for how drug screening assays should be performed and interpreted. Since many of the drug screening platforms work only at ambient temperature, measurements at different temperatures not only give rise to a large source of (deterministic) variation but also introduce the problem of translation of their findings to physiological temperatures. This translation is particularly problematic when an imperfect temperature model is used, such as the commonly used Q_10_ coefficient, as shown in this study. Extreme caution should be taken when using temperature-extrapolated *in vitro* drug screening data in *in silico* models for risk prediction.

Given we have shown Q_10_ coefficients definitely cannot capture the full temperature dependence of hERG kinetics (as shown in Figure 7), and different drugs target different kinetics, then previous findings that there are no common set of Q_10_ coefficients to describe the kinetics of drug block (52), are consistent with our results. In future, one could use our models to study some of the temperature effects observed in drug studies (11).

## CONCLUSION

We have studied the temperature dependence of hERG kinetics using a 15 second high-information content protocol developed in Part I of this study (16). We characterised the temperature dependence by fitting a mathematical model of hERG channel kinetics to data obtained at five distinct temperatures between 25 and 37 °C. We constructed between 45 and 124 cell-specific hERG models at each temperature using the 15 second calibration protocol, and our cell-specific variants of the hERG model were able to predict currents under eight independent validation protocols with high accuracy. We represented the variability in parameters using a hierarchical Bayesian model, and were able to reproduce the temperature dependence observed in previous literature studies. Our hERG models reveal that, overall, the activation process has a higher temperature sensitivity than the inactivation process. The temperature dependence of the kinetic parameters we obtained takes a more complicated form than that predicted by Q_10_ coefficients or a Typical Eyring approach, although it broadly follows a Generalised Eyring relationship. Our results show that a direct fit to the 15 second protocol is the best representation of hERG kinetics at a given temperature, although predictions from the Generalised Eyring theory may be preferentially used if no such data are available.

## AUTHOR CONTRIBUTIONS

CLL, MC, KAB, DJG, LP, KW & GRM designed the research. CLL, DM, JCH, KAB, LP & KW carried out pilot studies and the experiments shown here. CLL, MC, DJG & GRM designed the computational analysis. CLL wrote simulation codes, performed the analysis, and generated the results figures. All authors wrote and approved the final version of the manuscript.

## ACKNOWLEDGEMENTS

This work was supported by the Wellcome Trust [grant numbers 101222/Z/13/Z and 212203/Z/18/Z]; the Engineering and Physical Sciences Research Council and the Medical Research Council [grant number EP/L016044/1]; and the Biotechnology and Biological Sciences Research Council [grant number BB/P010008/1]. CLL acknowledges support from the Clarendon Scholarship Fund; and the EPSRC, MRC and F.Hoffman-LaRoche Ltd for studentship support via the Oxford Systems Approaches to Biomedical Science Centre for Doctoral Training. MC and DJG acknowledge support from a BBSRC project grant. GRM acknowledges support from the Wellcome Trust & Royal Society via a Sir Henry Dale Fellowship and a Wellcome Trust Senior Research Fellowship.

LP and KW are employees of F.Hoffman-LaRoche Ltd. and KW is a shareholder. KAB is an employee and shareholder of GlaxoSmithKline Plc.

## SUPPORTING CITATIONS

Reference (53) appears in the Supporting Material.

## SUPPORTING MATERIAL

An online supplement accompanies this article. All codes and data are freely available at https://github.com/CardiacModelling/hERGRapidCharacterisation.

