## Supporting Material for "Rapid characterisation of hERG channel kinetics II: temperature dependence"

### Supporting Material for:
### Rapid characterisation of hERG channel kinetics II

**Chon Lok Lei[1], Michael Clerx[1], Kylie A. Beattie[2], Dario Melgari[3], Jules C. Hancox[3], David J. Gavaghan[1], Liudmila Polonchuk[4], Ken Wang[4], and Gary R. Mirams[5]**

[1]Computational Biology, Department of Computer Science, University of Oxford, Oxford, United Kingdom
[2]Systems Modeling and Translational Biology, GlaxoSmithKline R&D, Stevenage, UK
[3]School of Physiology, Pharmacology and Neuroscience, and Cardiovascular Research Laboratories, School of Medical Sciences, University of Bristol, Bristol, United Kingdom
[4]Pharma Research and Early Development, Innovation Center Basel, F. Hoffmann-La Roche Ltd., Basel, Switzerland
[5]Centre for Mathematical Medicine and Biology, School of Mathematical Sciences, University of Nottingham, Nottingham, United Kingdom

#### Contents

**S1 Electrophysiology solutions** **1**
**S2 Recording techniques** **1**
**S3 Maximum conductance Estimation** **2**
**S4 Temperature-dependent fits and predictions** **2**
**S5 Relative root mean square error (RRMSE) histograms** **7**
**S6 Temperature dependence of inferred model parameters** **11**
**S7 Other methods for temperature-dependent models** **11**
**S8 Simulating literature temperature dependence** **12**
**S9 Simulating literature $Q_{10}$ estimation** **14**
**References** **15**

#### S1 Electrophysiology solutions

The compositions of all the electrophysiology solutions, including both the external solutions (bath solutions) and the internal solution (equivalent to the pipette solution in manual patch clamp), are shown in Table S1. External solutions were added in the following order: first 'fill chip' solution to the measurement chip, and the suspended hERG cells, then the 'seal enhancer' solution for enhancing the seal by forming CaF crystal around the cells (note they have extra high concentration of $Ca^{+}$, so we need to reduce/dilute it later), followed by adding the extracellular 'reference' solution for $Ca^{+}$ dilution. All the voltage clamp measurements were performed after adding all these external solutions.

The solutions were added sequentially to the wells, by removing half of the previous solutions from the wells each time. Therefore, the final ratios of the external (extracellular) solution are 1:1:2 — proportions of 0.25 of the 'Fill Chip' concentrations, 0.25 of the 'Seal Enhancer' concentrations, and 0.5 of the 'Reference' concentrations, as shown in the 'Final Extracellular' solution in Table S1.

#### S2 Recording techniques

All experiments were performed with Nanion SyncroPatch 384PE machine with software PatchControl384PE (v. 1.5.6 Build 22) and current traces data were exported using their complementary software DataControl384 (v. 1.5.0 Customer Release). The

| Solution<br>pH value (titrated with)<br>Osmolarity [mOsm] | | *Intracellular*<br>*pH 7.2 (KOH)*<br>*260-300* | *Fill Chip*<br>*pH 7.4 (NaOH)*<br>*300-330* | *Seal Enhancer*<br>*pH 7.4 (HCl)*<br>*290-330* | *Reference*<br>*pH 7.4 (HCl)*<br>*290-330* | *Final Extracellular* |
|---|---|---|---|---|---|---|
| **Chemicals** | **Source / Cat#** | **[ ] in mM** | **[ ] in mM** | **[ ] in mM** | **[ ] in mM** | **[ ] in mM** |
| NaCl | Merck / K38447104807 | 10 | 150 | 80 | 80 | 97.5 |
| KCl | Merck / K36782536 | 10 | 4 | 4 | 4 | 4 |
| KF | Acros Organics / 201352500 | 100 | — | — | — | — |
| $MgCl_2$ | Merck / A914133908 | — | 1 | 1 | 1 | 1 |
| $CaCl_2$ | Acros Organics/ 349615000 | — | 1.2 | 5 | 1 | 2.05 |
| HEPES | Applichem A1069 | 10 | 10 | 10 | 10 | 10 |
| Glucose | Fluka / 49159 | — | 5 | 5 | 5 | 5 |
| NMDG | Fluka 66930 | — | — | 60 | 40 | 35 |
| EGTA | Fluka / 03778 | 20 | — | — | — | — |
| Sorbitol | Sigma / S1876 | — | — | — | 40 | 20 |

**Table S1.** Electrophysiology solutions for hERG assay on the Nanion SyncroPatch 384PE machine, all solutions are sterile filtered. All hERG cells were suspended in 1/3 Extracellular Fill Chip Solution + 2/3 Hanks' Balanced Salt Solution (HBSS).

machine comes with a measurement chip consists of 364 wells, with 16 rows by 24 columns.

#### S3 Maximum conductance Estimation

Figure S1 shows an illustration of the estimation of maximum conductance value. The estimation is done by extrapolating the negative tail current (blue), after the first 40 mV to $-120$ mV step, back to the time the voltage step occurred (green vertical dashed line). The extrapolation is done by fitting a single exponential function (orange dashed line) to the tail current. The estimated value is indicated by the black cross. The absolute value is used as the normalisation constant of the current. Note that this value is only used for normalising plots and comparing between cells/temperatures (as the channel is close to fully active ($a \approx 1$) at this time point at all temperatures). The value is *not* used in the mathematical model fitting/validation for an individual well.

#### S4 Temperature-dependent fits and predictions

Figures S2, S3 and S4 show the model fitting and validation results for all recorded cells at 27, 30, and 33 °C respectively, under the 9 different protocols. From (panel A) to (panel I): the staircase protocol, the activation current-voltage (I-V) protocol, the steady-state inactivation I-V protocol, the hERG screening protocol, the DAD-like protocol, the EAD-like protocol, and the cardiac action potential-like protocol at 0.5 Hz, 1 Hz and 2 Hz, respectively. All model predictions are compared against the experimental recordings measured under the same protocols. We fitted the model to the staircase protocol (panel A) and validated against the other 8 protocols (panels B–I). To compare the variability in hERG kinetics only, currents are normalised[1] by scaling them to minimise the absolute difference between each trace and a reference trace.

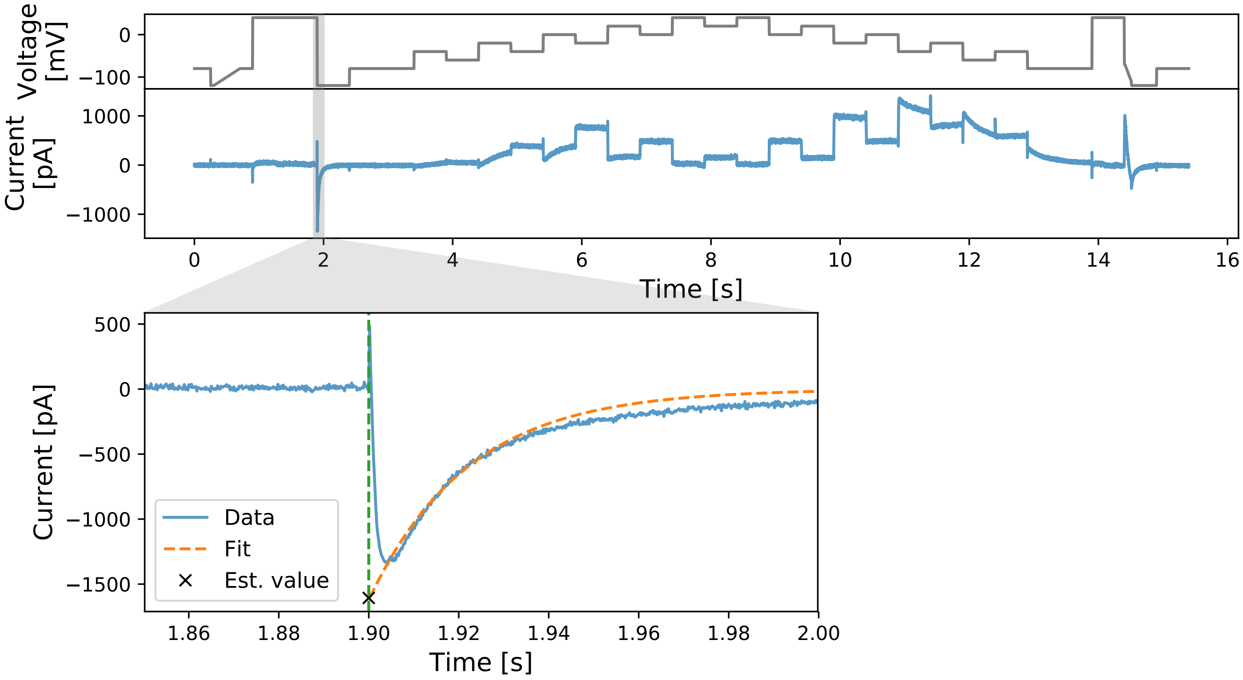


**Figure S1.** An illustration of the estimation of the maximum conductance value (black cross). In the magnified plot, the negative tail current (blue) after the first 40 mV to $-120$ mV step is extrapolated back to the time the voltage step occurred (indicated by the green vertical dashed line). The extrapolation is done by fitting a single exponential function (orange dashed line) to the tail current.

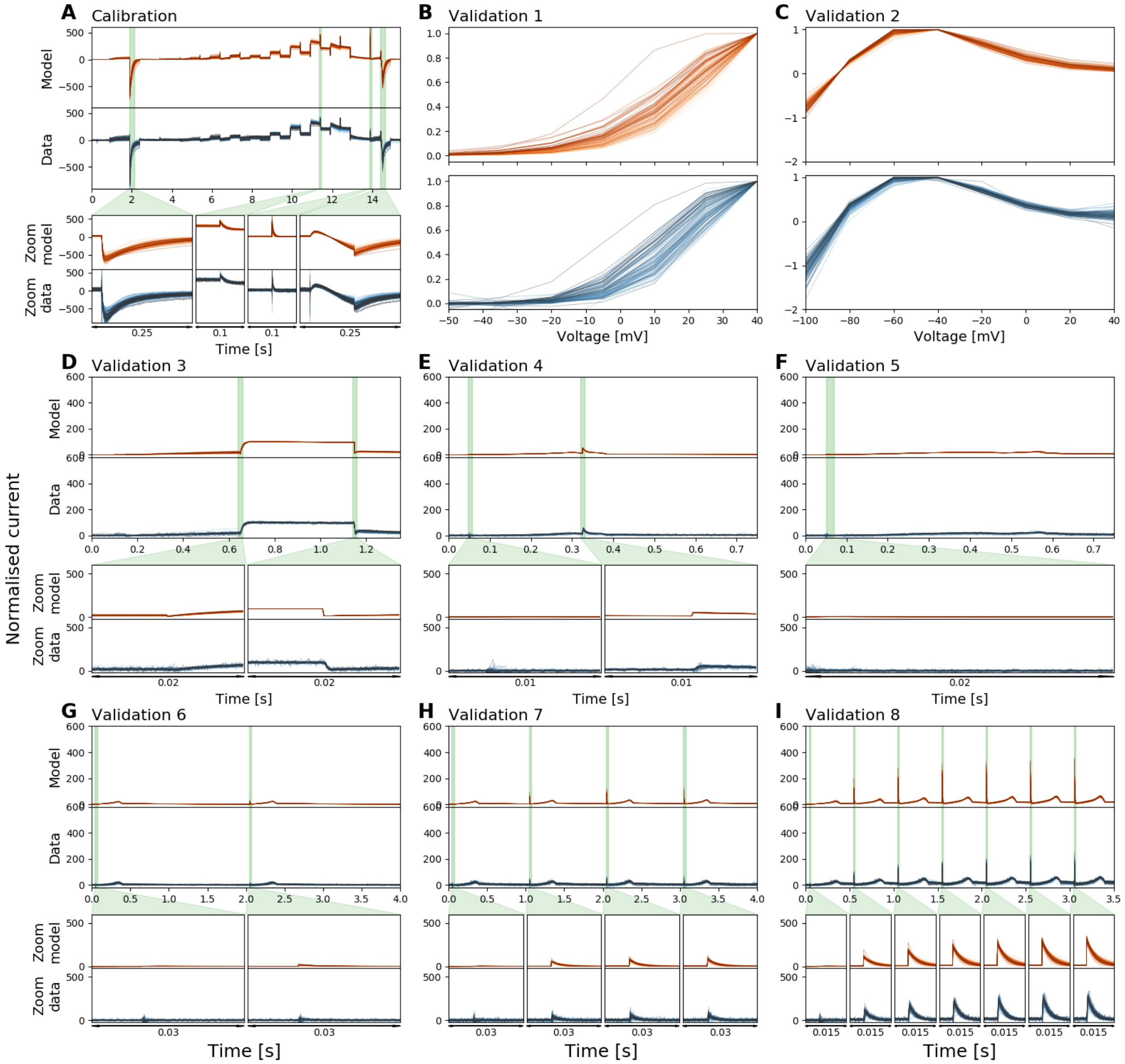


**Figure S2.** Normalised whole-cell patch-clamp voltage clamp recordings under 9 different protocols, and the model fitting and validation results, at 27 °C. All currents are normalised by scaling them to minimise the absolute difference between each trace and a reference trace. From **(A)** to **(I)**: The staircase protocol which is used as the calibration protocol, the activation current-voltage (I-V) protocol, the steady-state inactivation I-V protocol, the hERG screening protocol, the DAD-like protocol, the EAD-like protocol, and the cardiac action potential-like protocol at 0.5 Hz, 1 Hz and 2 Hz, respectively. All the model calibration results and validation predictions are shown in the top panels (orange), and are compared against the experimental recordings shown in the bottom panels (blue). Zoomed-in of the green shaded regions are shown underneath each panel to reveal the details of the spikes, in which our models show extraordinary good predictions to the details. The normalised current for all protocols are shown except for the activation I-V protocol and the steady-state inactivation I-V protocol where the summary statistic I-V relationships are shown. Each cell is shown with a unique colour.

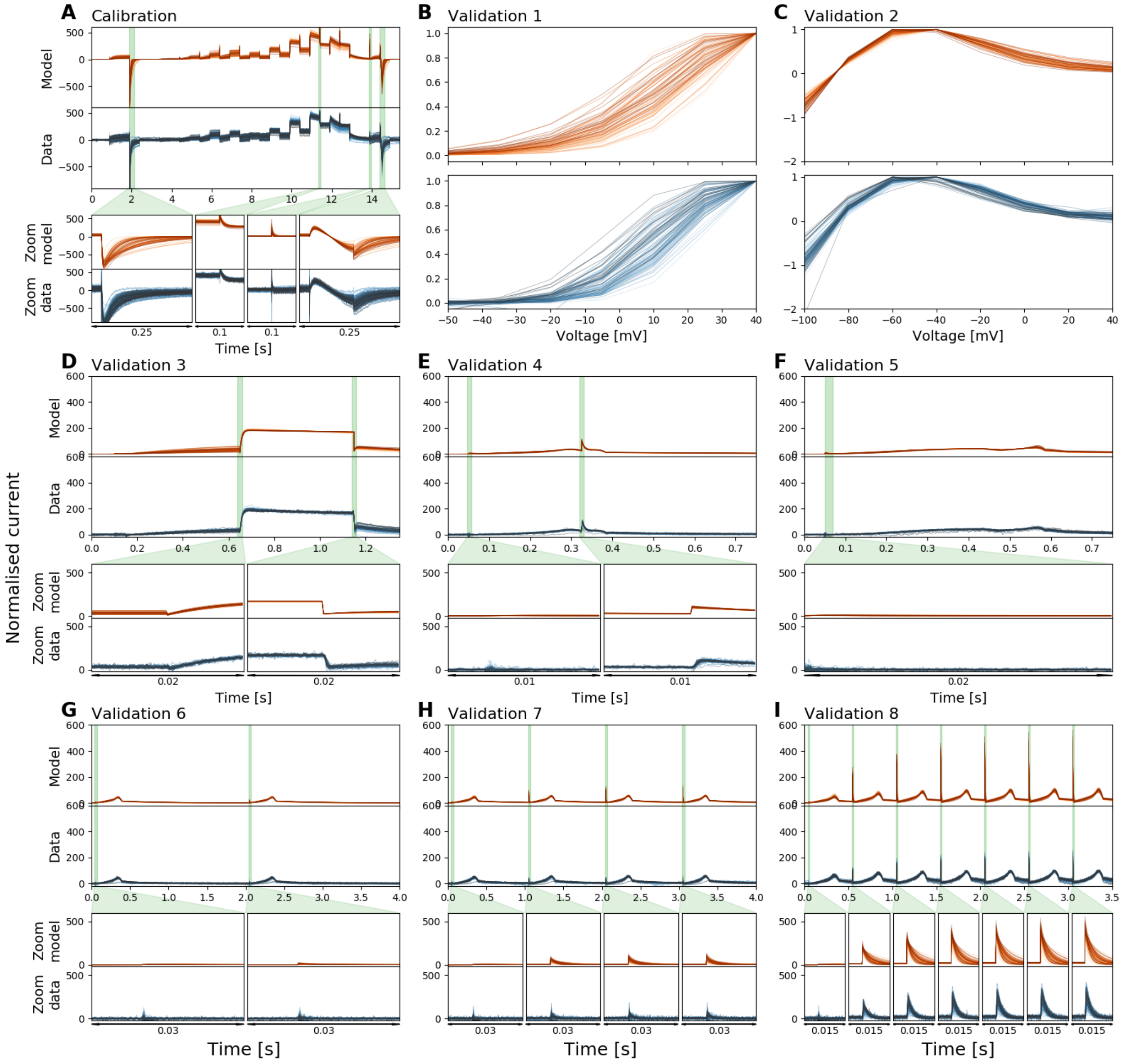


**Figure S3.** Normalised whole-cell patch-clamp voltage clamp recordings under 9 different protocols, and the model fitting and validation results, at 30 °C. All currents are normalised by scaling them to minimise the absolute difference between each trace and a reference trace. From **(A)** to **(I)**: The staircase protocol which is used as the calibration protocol, the activation current-voltage (I-V) protocol, the steady-state inactivation I-V protocol, the hERG screening protocol, the DAD-like protocol, the EAD-like protocol, and the cardiac action potential-like protocol at 0.5 Hz, 1 Hz and 2 Hz, respectively. All the model calibration results and validation predictions are shown in the top panels (orange), and are compared against the experimental recordings shown in the bottom panels (blue). Zoomed-in of the green shaded regions are shown underneath each panel to reveal the details of the spikes, in which our models show extraordinary good predictions to the details. The normalised current for all protocols are shown except for the activation I-V protocol and the steady-state inactivation I-V protocol where the summary statistic I-V relationships are shown. Each cell is shown with a unique colour.

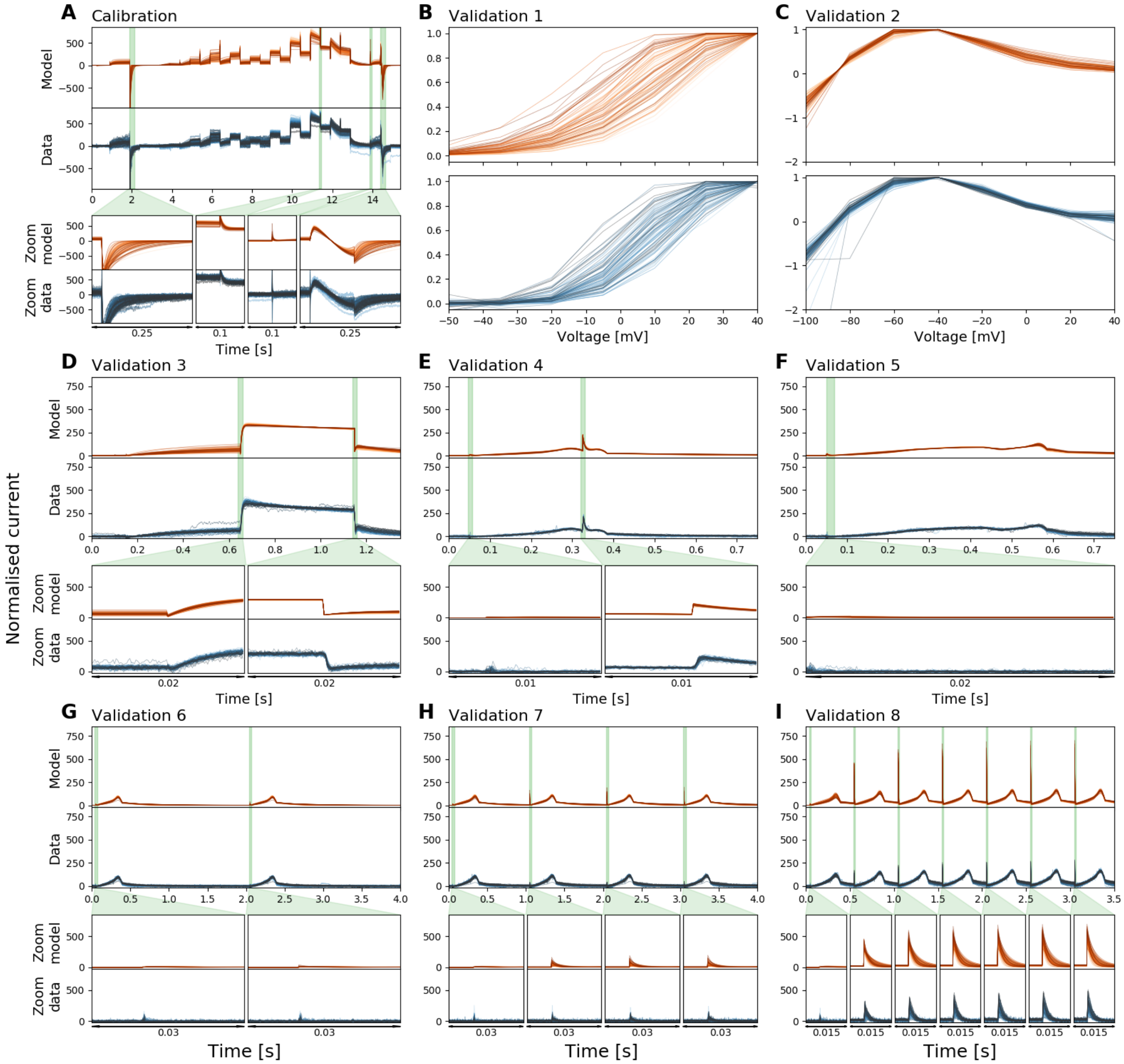


**Figure S4.** Normalised whole-cell patch-clamp voltage clamp recordings under 9 different protocols, and the model fitting and validation results, at 33 °C. All currents are normalised by scaling them to minimise the absolute difference between each trace and a reference trace. From **(A)** to **(I)**: The staircase protocol which is used as the calibration protocol, the activation current-voltage (I-V) protocol, the steady-state inactivation I-V protocol, the hERG screening protocol, the DAD-like protocol, the EAD-like protocol, and the cardiac action potential-like protocol at 0.5 Hz, 1 Hz and 2 Hz, respectively. All the model calibration results and validation predictions are shown in the top panels (orange), and are compared against the experimental recordings shown in the bottom panels (blue). Zoomed-in of the green shaded regions are shown underneath each panel to reveal the details of the spikes, in which our models show extraordinary good predictions to the details. The normalised current for all protocols are shown except for the activation I-V protocol and the steady-state inactivation I-V protocol where the summary statistic I-V relationships are shown. Each cell is shown with a unique colour.

#### S5 Relative root mean square error (RRMSE) histograms

The relative root mean square error (RRMSE, defined in Eq. 25 in the main text) analysis and the resulting histograms for 27, 30, and 33 °C are shown in Figures S5, S6, and S7 respectively. All the results demonstrate that our hERG models are good representations of the kinetics of the cells at each corresponding temperature.

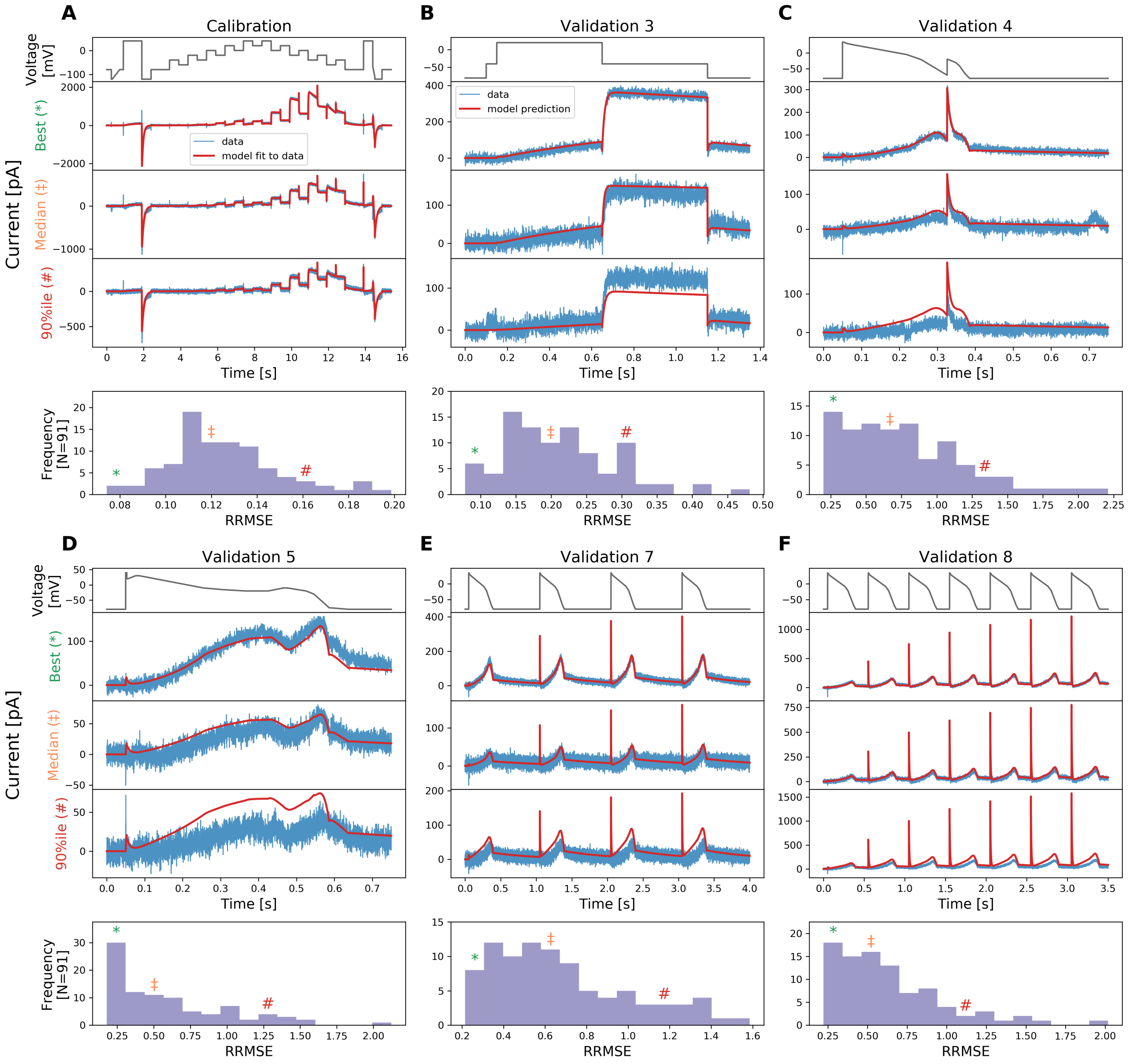


**Figure S5.** The relative root mean square error (RRMSE, given by Eq. 25) histograms for all cells and for 6 protocols used at 27 °C. Markers indicate the best (∗), median (‡) and 90$^{th}$ percentile (#) RRMSE values. The raw traces with the best, median and 90$^{th}$ percentile RRMSE values, for both the model (red) and data (blue), are shown in the panels above, together with the voltage protocol shown on top. Note that the currents are shown on different $y$-axis limits, to reveal the details of the traces.

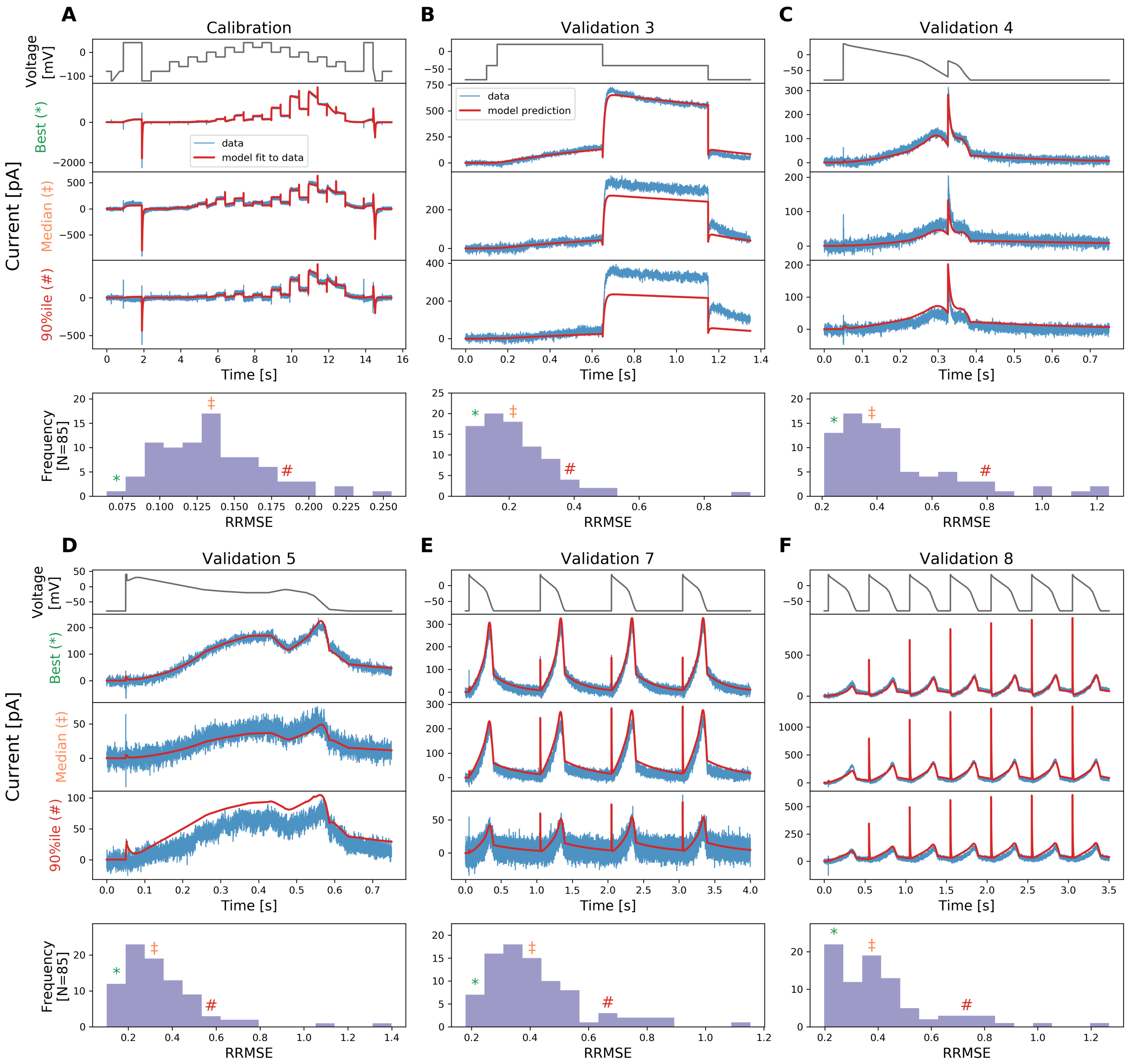


**Figure S6.** The relative root mean square error (RRMSE, given by Eq. 25) histograms for all cells and for 6 protocols used at 30 °C. Markers indicate the best ($*$), median (‡) and 90$^{\text{th}}$ percentile (#) RRMSE values. The raw traces with the best, median and 90$^{\text{th}}$ percentile RRMSE values, for both the model (red) and data (blue), are shown in the panels above, together with the voltage protocol shown on top. Note that the currents are shown on different $y$-axis limits, to reveal the details of the traces.

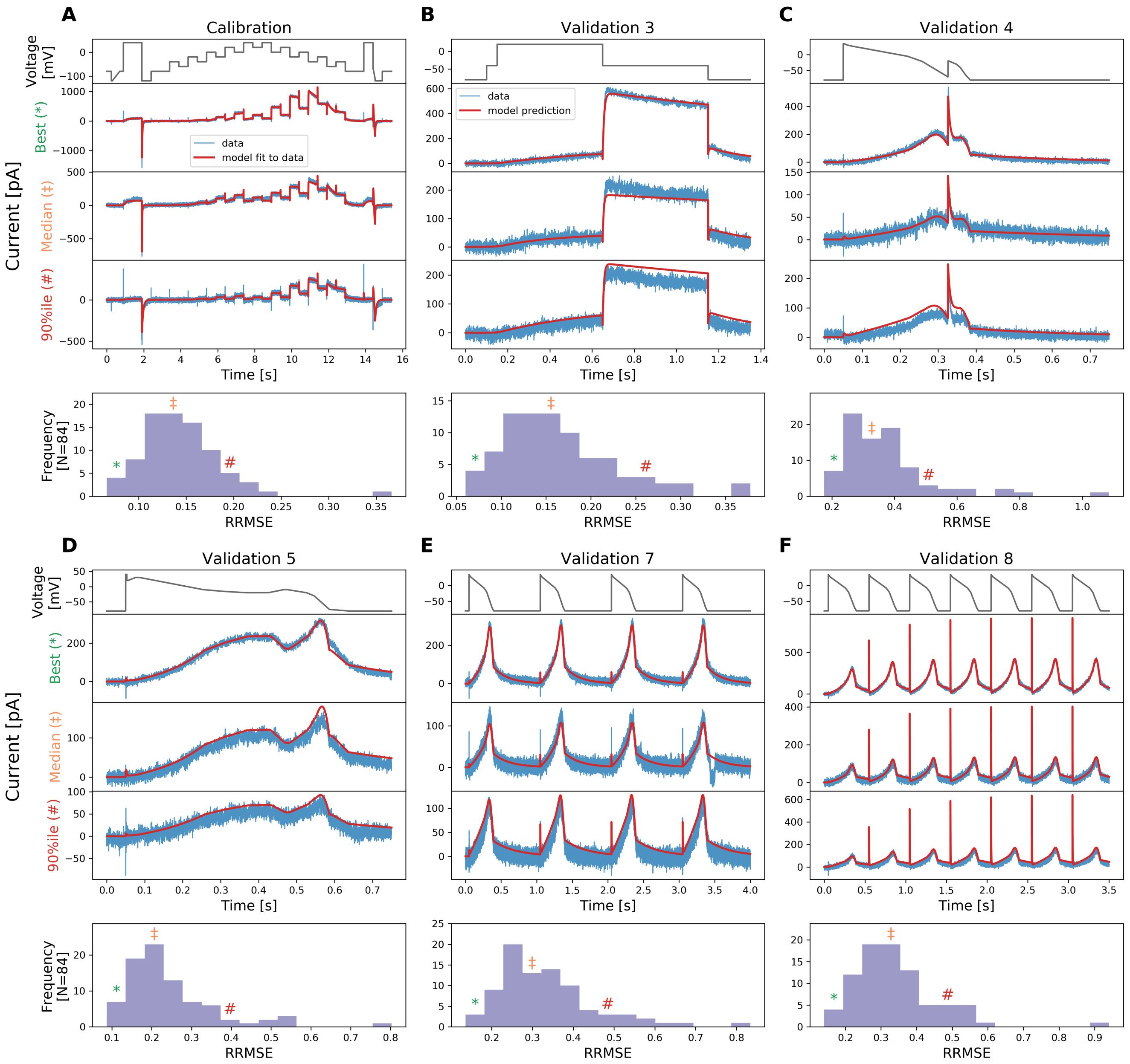


**Figure S7.** The relative root mean square error (RRMSE, given by Eq. 25) histograms for all cells and for 6 protocols used at 33 °C. Markers indicate the best (∗), median (‡) and 90$^{\text{th}}$ percentile (#) RRMSE values. The raw traces with the best, median and 90$^{\text{th}}$ percentile RRMSE values, for both the model (red) and data (blue), are shown in the panels above, together with the voltage protocol shown on top. Note that the currents are shown on different $y$-axis limits, to reveal the details of the traces.

#### S6 Temperature dependence of inferred model parameters

Figure S8 shows the inferred parameter values as a function of temperature. The results are shown in the Eyring plot form, where $\ln(A/T)$ and $B$ are plotted against $T^{-1}$. It shows the inferred distribution of the hyperparameter mean $\mu$ (Eq. 24 in the main text) using the simplified pseudo-MwG at each temperature in a violin plot. In Figure S8, parameters $p_1, p_3, p_4, p_5, p_7, p_8$ show a linear trend as temperature increases, as predicted by Figure 1 in the main text; whereas parameters $p_2, p_6$ take a slightly more complicated form.

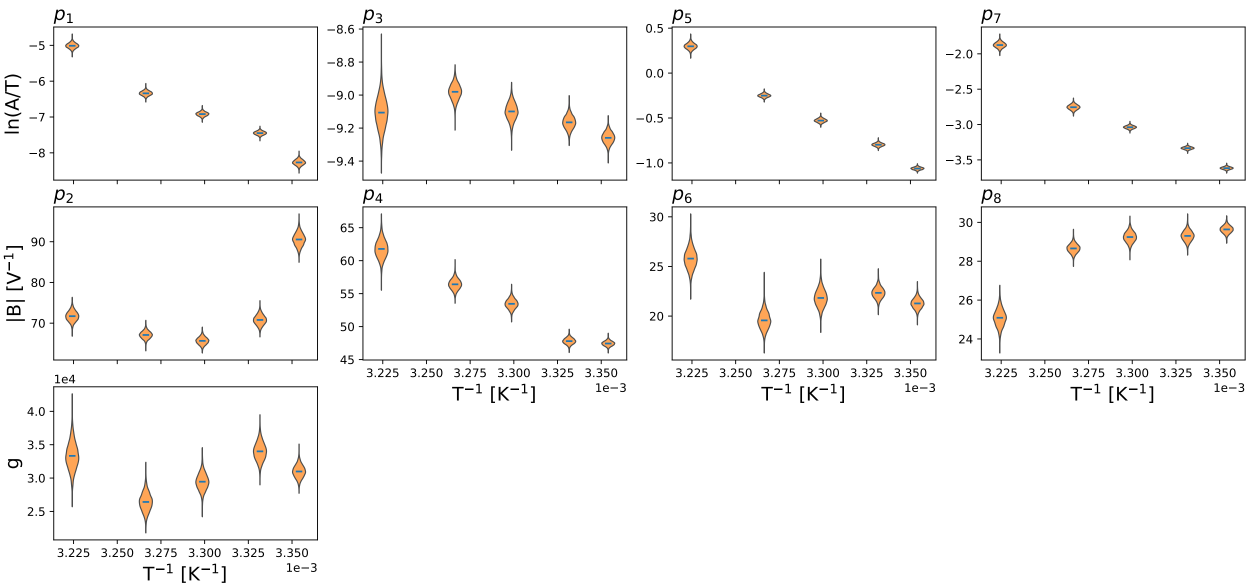


**Figure S8.** Model parameters plotted as a function of temperature in an Eyring plot: $\ln(A/T)$ and $B$ are plotted against $T^{-1}$. Here only the inferred distribution of the hyperparameter mean $\mu$ (Eq. 24 in the main text) using the simplified pseudo-MwG at each temperature is shown. Model parameters show different degrees of temperature dependency. The conductance $g$ does not show a prominent change as temperature increases.

#### S7 Other methods for temperature-dependent models

Figure S9 shows the fitted Generalised Eyring relationship and the $Q_{10}$ equation to the inferred parameters shown in Figure S8 (orange violin plot). The obtained Generalised Eyring fits are shown as green fan charts with the first three standard deviations in green; the obtained $Q_{10}$ fits are shown in red. The fitted parameters for the Generalised Eyring equation (Eq. 4 in the main text) and the $Q_{10}$ equation (Eq. 9 in the main text) are given in the bottom right tables, one set for each $k_i$, $i = 1, 2, 3, 4$.

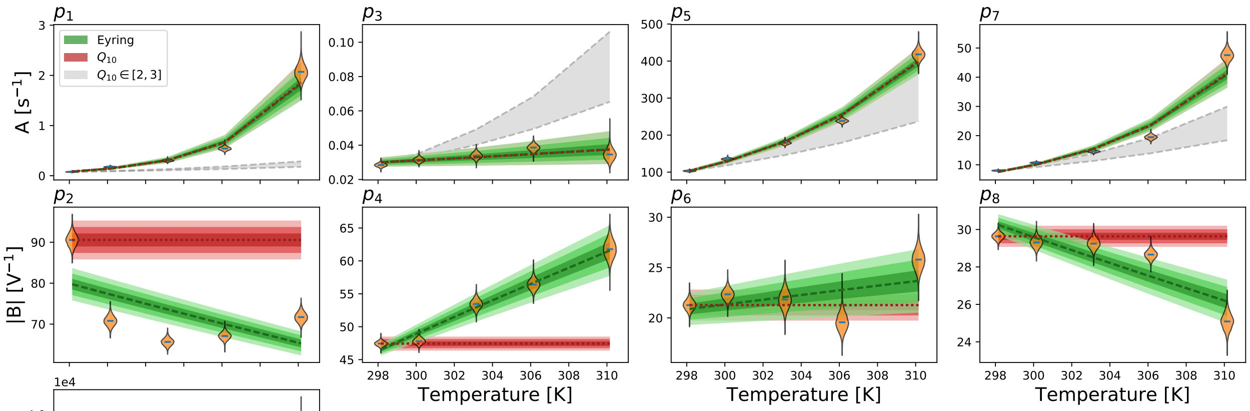


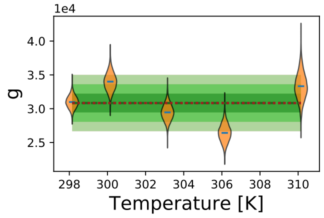


| | $\Delta S$ (JK$^{-1}$mol$^{-1}$) | $\Delta H$ (Jmol$^{-1}$) | $z_e$ | $D$ (V$^{-1}$) |
|---|---|---|---|---|
| $k_1$ | 389.543 | 195330.0 | 9.59193 | -293.601 |
| $k_2$ | -235.457 | 11514.1 | 9.98987 | -435.358 |
| $k_3$ | 75.3932 | 84017.4 | -1.85611 | 93.1319 |
| $k_4$ | 125.715 | 105487.0 | -2.71481 | 75.4196 |

| | $Q_{10}$ | $\alpha$ (s$^{-1}$) | $\beta$ (V$^{-1}$) |
|---|---|---|---|
| $k_1$ | 13.1283 | 0.0843731 | 90.5512 |
| $k_2$ | 1.19849 | 0.030066 | -47.4353 |
| $k_3$ | 3.0847 | 103.436 | 21.2742 |
| $k_4$ | 4.08594 | 7.62406 | -29.6366 |

**Figure S9.** Fitting of Generalised Eyring equation and $Q_{10}$ equation to the mean distribution $\mu$ inferred using the simplified psuedo-MwG (orange violin plot). The obtained Generalised Eyring fits are shown as green fan charts with the first three standard deviations in green; the obtained $Q_{10}$ fits are shown in red. The fitted parameters for the Generalised Eyring and $Q_{10}$ equations are shown in the bottom right tables, one set for each $k_i$, $i = 1, 2, 3, 4$. For comparison to typical $Q_{10}$ values in literature, where $Q_{10}$ values are usually assumed to be around 2 to 3, we show the parameters prediction using $Q_{10} \in [2, 3]$ as the grey shaded region.

#### S8 Simulating literature temperature dependence

Here we describe the procedure for reproducing the literature results of Vandenberg *et al.*[2] (Figure 6) through model simulations. These authors estimated the 'open probability' through experimental measurements. We use our directly fitted hierarchical Bayesian models at 25 °C and 33 °C for simulation, and compare to their reported temperature induced changes with measurements performed at 22 °C and 32 °C. The results are shown in Figure 10 and are discussed in the Discussion section in the main text.

First, we simulate the voltage dependence of activation using the 'isochronal tail current protocol'[2]: a holding potential of −80 mV, followed by 30 s depolarising pulses to voltages from −80 to 40 mV, then 500 ms steps to −120 mV. The tail currents during the −120 mV steps are recorded to construct the voltage dependence of activation curves shown in Figure S10. Our results here are compared to Figure 3C in Vandenberg *et al.*[2].

Second, we simulate the steady-state inactivation using the 'double-pulse protocol'[2]: a holding potential of −80 mV, followed by 1 s depolarising pulses to voltage 40 mV, then 500 ms steps to voltages in the range −120 to 40 mV. The peak amplitude of the tail current is corrected using the method as described in Section S3, the same method used in Vandenberg *et al.*[2]. These corrected current values are then converted to conductances by dividing by the electrochemical driving force $(V - E_K)$. These are described as the steady-state inactivation curves[2] and are shown in Figure S11. Our results here are compared to Figure 5D in Vandenberg *et al.*[2].

The product of the voltage dependence of activation (Figure S10) and the steady-state inactivation (Figure S11) gives the 'open probability' defined in Vandenberg *et al.*[2] (Figure 6), which is shown in Figure 10 in the main text. Strictly speaking, this 'open probability' is *not* the open probability of the model ($O = a \cdot r$), but an approximate of $O$ from the simulation results following simulation of the experimental protocols and a repeat of the analyses in Vandenberg *et al.*[2] based on the simulated currents.

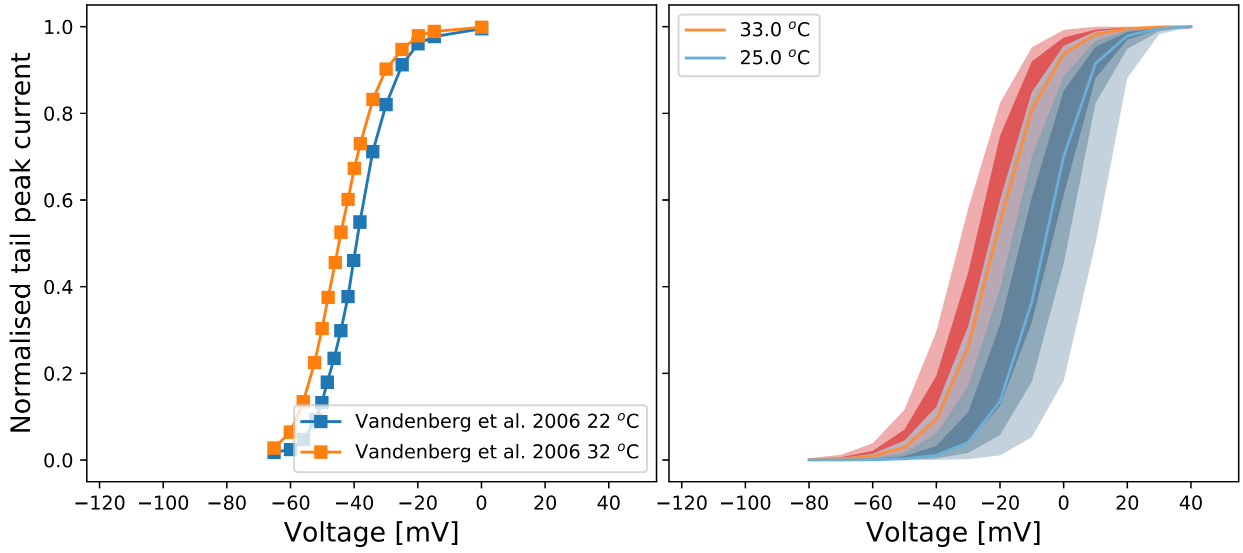


**Figure S10.** Voltage dependence of activation defined in Vandenberg *et al.*[2] (Figure 3C). **Left:** Data reproduced from Vandenberg *et al.*[2] (Figure 3C). **Right:** The fan charts show the the 90$^{th}$, 60$^{th}$ and 30$^{th}$ percentiles of the hierarchical Bayesian model simulations, representing the *experiment-experiment* variability. Orange/red represents 32 °C to 33 °C, and blue represents 22 °C to 25 °C.

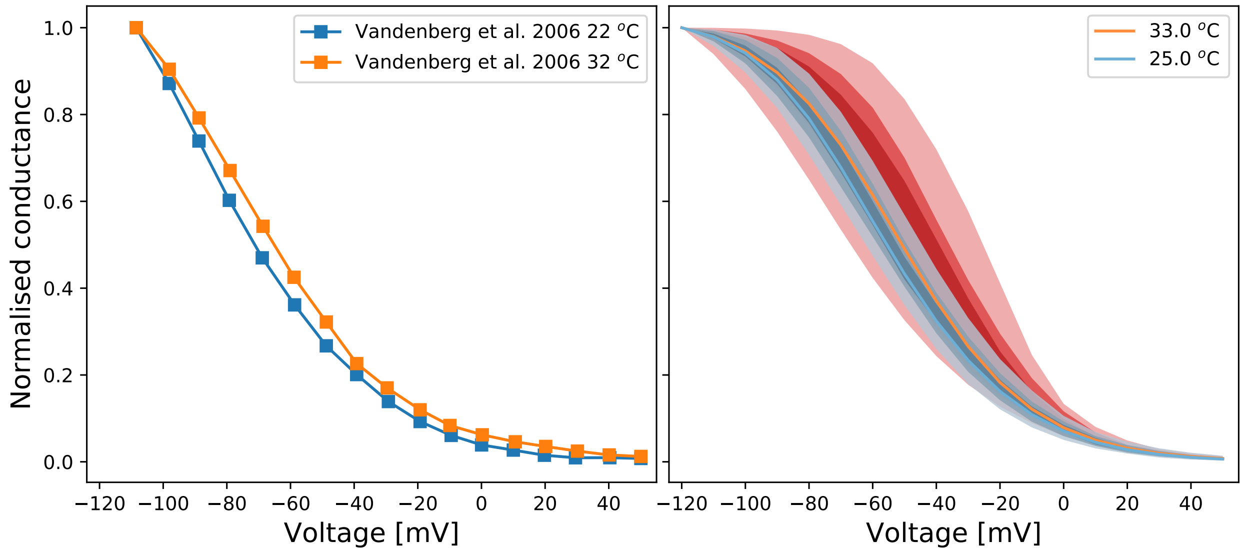


**Figure S11.** Steady-state inactivation defined in Vandenberg *et al.*[2] (Figure 5D). **Left:** Data reproduced from Vandenberg *et al.*[2] (Figure 5D). **Right:** The fan charts show the the 90$^{th}$, 60$^{th}$ and 30$^{th}$ percentiles of the hierarchical Bayesian model simulations, representing the *experiment-experiment* variability. Orange/red represents 32 °C to 33 °C, and blue represents 22 °C to 25 °C.

We also compare the temperature effect under action potential clamps. In Figure S12, we plot experimental data from our high-throughput measurements on the left, and compare with the simulations under the protocol shown in Figure 1B of Vandenberg *et al.*[2]. These figures show similar temperature effects due to increasing temperature from 25 °C (blue) to 33 °C (red). The simulation on the right is comparable to that in Figure 1B of Vandenberg *et al.*[2]. We are able to reproduce the results that are broadly consistent with Vandenberg *et al.*[2].

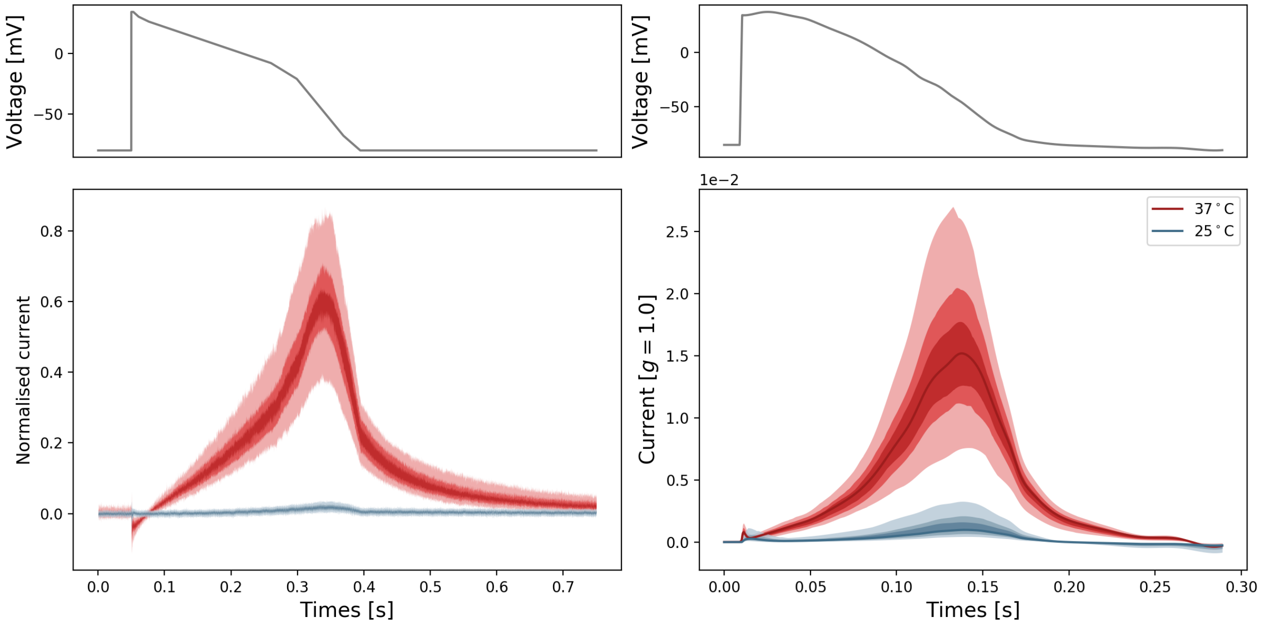


**Figure S12.** Action potential clamp (left) data from our high-throughput measurements and (Right) simulations under the protocol in Vandenberg *et al.*[2] (Figure 1B). The fan charts show the the 30$^{\text{th}}$, 60$^{\text{th}}$ and 90$^{\text{th}}$ percentiles of the data and the hierarchical Bayesian model simulations, representing the *experiment-experiment* variability. Orange/red represents 32 °C to 33 °C, and blue represents 22 °C to 25 °C.

#### S9 Simulating literature $Q_{10}$ estimation

Here we describe the procedure for reproducing the $Q_{10}$ coefficients reported by Zhou *et al.*[3] and Vandenberg *et al.*[2] through model simulations. We use our directly fitted hierarchical Bayesian models at 25 °C and 37 °C to compute $Q_{10}$ coefficients by simulating the experimental procedure that was used to derived the literature $Q_{10}$ coefficients. The obtained $Q_{10}$ are shown in Table 1 and are discussed in the Discussion section in the main text.

##### $Q_{10}$ coefficients in Zhou *et al.*[3]

We simulate a depolarising step from −80 mV (holding potential) to 0 mV for 5 s which followed by a repolarising step to −50 mV for 5 s. The time constant, fitted with a single exponential, of the current during 0 mV is called the activation time constant. The fitted time constant of the tail current during the −50 mV step is called the deactivation time constant.

We then simulate the ‘three-pulse protocol’[3]: a holding potential of −80 mV, a 200 ms depolarising step to 60 mV followed by a repolarising step to −100 mV for 2 ms, then a step to voltage 0 mV for 200 ms. The fitted time constant of the current during 0 mV is called the inactivation time constant. We finally simulate the two-pulse protocol[3]: a holding potential of −80 mV, followed by a 200 ms depolarising step to 60 mV before a repolarising step to −50 mV for 200 ms. The fitted time constant of the tail current rising phase during −50 mV is called the recovery (from inactivation) time constant.

#### $Q_{10}$ coefficients in Vandenberg *et al.*[2]

We simulate the 'envelope of tails protocol'[2]: a holding potential of −80 mV, followed by a depolarising step to 0 mV for variable durations before stepping voltage to −50 mV for 500 ms. The peak amplitudes of the tail currents during the −50 mV steps are plotted against the duration of the 0 mV step. The time constant, fitted with a single exponential function, of the peak current-step duration curve is called the activation time constant.

We then simulate the 'triple pulse protocol'[2]: a holding potential of −80 mV, a 1 s depolarising step to 40 mV followed by a repolarising step to −80 mV for 20 ms, then a step to voltage 40 mV for 100 ms, and finally a step to voltage −120 mV for 200 ms. The fitted time constant of the current during the second 40 mV is called the inactivation time constant. Two time constants $\tau_1, \tau_2$ are obtained by fitting a 'double exponential' function

$$f_{\text{double exponential}}(t) = A\exp(-t/\tau_1) - B\exp(-t/\tau_2), \tag{S1}$$

with constants $A, B$, to the tail current during −120 mV. The time constant corresponding to the initial increase in inward current is called the recovery (from inactivation) time constant; the time constant of the decrease in inward current is called the deactivation time constant.

#### Computing mean and standard deviation of $Q_{10}$ coefficients

We compare the results using mean and standard deviation, such that we can take the variability of experiments into account. Instead of the time constant, we use its inverse, the rate constant $\kappa$, for further calculation. For each type of rate constant (activation, deactivation, inactivation, and recovery), we compute the mean $\hat{\kappa}$ and standard deviation $\delta\kappa$ using all the simulated results for each temperature.

Since $Q_{10}$ is given by

$$Q_{10} = \left(\frac{\kappa_1}{\kappa_2}\right)^{10°\mathrm{C}/(T_2-T_1)}, \tag{S2}$$

where $\kappa_1, \kappa_2$ are rate constants at temperature $T_1 = 25\,°\mathrm{C}$, $T_2 = 37\,°\mathrm{C}$ respectively. We assume a linear error and independent variables, and denote the exponent $10°\mathrm{C}/(T_2 - T_1)$ as $\Delta T$, the standard deviation of a variable $x$ as $\delta x$, and its mean estimator as $\hat{x}$. Then we apply the propagation of error equation[4] which gives

$$\delta Q_{10} = \left\{ \delta\kappa_1^2 \left(\frac{\partial Q_{10}}{\partial \kappa_1}\bigg|_{\hat{Q}_{10}}\right)^2 + \delta\kappa_2^2 \left(\frac{\partial Q_{10}}{\partial \kappa_2}\bigg|_{\hat{Q}_{10}}\right)^2 + \delta T_1^2 \left(\frac{\partial Q_{10}}{\partial T_1}\bigg|_{\hat{Q}_{10}}\right)^2 + \delta T_2^2 \left(\frac{\partial Q_{10}}{\partial T_2}\bigg|_{\hat{Q}_{10}}\right)^2 \right\}^{-1/2} \tag{S3}$$

$$= \left\{ \left(\delta\kappa_1 \frac{\hat{\Delta T}}{\hat{\kappa}_1}\hat{Q}_{10}\right)^2 + \left(\delta\kappa_2 \frac{\hat{\Delta T}}{\hat{\kappa}_2}\hat{Q}_{10}\right)^2 + \left(\delta T_1 \hat{Q}_{10} \ln\frac{\hat{\kappa}_1}{\hat{\kappa}_2}\frac{\hat{\Delta T}^2}{10°\mathrm{C}}\right)^2 + \left(\delta T_2 \hat{Q}_{10} \ln\frac{\hat{\kappa}_1}{\hat{\kappa}_2}\frac{\hat{\Delta T}^2}{10°\mathrm{C}}\right)^2 \right\}^{-1/2}, \tag{S4}$$

where we assume $\delta T \approx 1\,°\mathrm{C}$.

Using the mean values $\hat{\kappa}_1, \hat{\kappa}_2$ and Eq. S4, we compute an estimation of the mean and standard deviation of $Q_{10}$ for each type of rate constant, for Zhou *et al.*[3] and Vandenberg *et al.*[2]. The results are shown in the "Model estimation" columns in Table 1 in the main text.

Since Zhou *et al.*[3] reported only the time constants of each type with mean and standard error of mean $\sigma_{\bar{x}}$ at two temperatures, we first estimate the mean and standard deviation for rate constants. We approximate the standard deviation with $\approx \sqrt{n}\sigma_{\bar{x}}$, where $n$ is the number of cells reported for each time constant. Then we apply Eq. S4 to propagate the errors in rate constants at two temperatures to estimate the error in their $Q_{10}$coefficients. Vandenberg *et al.*[2] reported the $Q_{10}$ coefficient with mean and standard error of mean for each gating process, we convert the standard error of mean to standard deviation for comparison. The results are shown in the "Reported values" columns in Table 1.
